# Extrinsic control of the early postnatal CA1 hippocampal circuits

**DOI:** 10.1101/2022.06.03.494656

**Authors:** Erwan Leprince, Robin F. Dard, Salomé Mortet, Caroline Filippi, Marie Giorgi-Kurz, Pierre-Pascal Lenck-Santini, Michel A. Picardo, Marco Bocchio, Agnès Baude, Rosa Cossart

## Abstract

The adult CA1 region of the hippocampus produces coordinated neuronal dynamics with minimal reliance on its extrinsic inputs. In contrast, the neonatal CA1 is tightly linked to externally-generated sensorimotor activity but the circuit mechanisms underlying early synchronous activity in CA1 remain unclear. Here, using a combination of *in vivo* and *ex vivo* circuit mapping, calcium imaging and electrophysiological recordings in mouse pups, we show that early dynamics in the ventro-intermediate CA1 are under the mixed influence of entorhinal (EC) and thalamic (VMT) inputs. Both VMT and EC can drive internally-generated synchronous events *ex vivo*. However, movement-related population bursts detected *in vivo* are exclusively driven by the EC. These differential effects on synchrony reflect the different intrahippocampal targets of these inputs. Hence, cortical and subcortical pathways act differently on the neonatal CA1, implying distinct contributions to the development of the hippocampal microcircuit and related cognitive maps.

## INTRODUCTION

Cortical neurons are spontaneously active in a coordinated manner as they mature and integrate into functional networks (Blankenship and Feller, 2009; Cossart and Garel, 2022; Martini et al., 2021; Molnár et al., 2020). Spontaneous activity contributes to the progressive tuning of cells and circuits to function by adjusting initial genetic programs to the statistics of the environment and the body (Cossart and Garel, 2022). Given this major developmental role of early spontaneous neuronal activity, it is essential to understand the circuit mechanisms supporting its generation. These involve the interaction between local intracortical circuits and extrinsic inputs. This interaction between local and external drives can be reflected in the co-existence of different types of network patterns supported by separate circuits (Allene et al., 2008; Siegel et al., 2012).

While the local and peripheral circuits involved in the generation of early spontaneous activity start to be well-elucidated in the developing sensory neocortex (Luhmann and Khazipov, 2017; Martini et al., 2021; Molnár et al., 2020), this is not yet the case for the hippocampus (Cossart and Khazipov, 2021). This stems from the combined difficulty to experimentally access this region with optical approaches allowing for local circuit dynamics mapping at cellular resolution, and to integrate the various external inputs it receives into a comprehensive framework. Within the hippocampus, the CA1 region is under the conjunctive influence of many extra-hippocampal brain regions, including the medial and lateral entorhinal cortices (EC) conveying spatial and non-spatial (e.g. sensory) information, as well as the nucleus reuniens from the higher order ventral midline thalamus (VMT), conveying information from the prefrontal cortex (Cassel et al., 2021; Ferraris et al., 2021; Vertes, 2015). *Ex vivo* early CA1 dynamics are dominated by so-called Giant Depolarizing Potentials (Ben-Ari et al., 1989), largely relying on local GABAergic circuits for their generation (Bocchio et al., 2020; Wester and McBain, 2016). *In vivo,* early CA1 dynamics have been mainly described with extracellular electrophysiological recordings. They are dominated by early sharp waves (eSPWs), recently described as bottom-up events involving activation of the EC during myoclonic movements and a major sink in the *stratum lacunosum moleculare (slm)* indicating this CA1 sublayer as a significant site of excitation (Valeeva et al., 2019). In addition to interactions within the hippocampal formation, the link between movements and eSPWs may also involve subcortical pathways such as projections from the nucleus reuniens from the ventral midline thalamus directly to CA1, given that these: (i) like the EC, also provide a source of glutamatergic excitation terminating onto the *slm*, (ii) develop prior to intrahippocampal circuits, (iii) have a mild, but significant functional impact on the occurrence of hippocampal theta bursts (Hartung et al., 2016), and convey important top-down information from the prefrontal cortex.

Here we have used a combination of *ex vivo* and *in vivo* circuit mapping with calcium imaging, optogenetics and chemogenetics to disentangle the respective contributions of EC and VMT inputs to the coordination of local CA1 dynamics at the end of the first postnatal week in mice. We show that both inputs can significantly modulate early CA1 dynamics recorded *ex vivo* and *in vivo* but differ in their distribution and preferred postsynaptic targets. While the EC most often directly triggers CA1 synchronization through large amplitude EPSCs onto CA1 pyramids, VMT inputs rather exert a modulatory indirect effect onto these cells. Still, the VMT can synchronize activity *ex vivo* through the almost exclusive activation of CA1 *slm* interneurons.

## MATERIALS AND METHODS

### MICE

All experiments were performed under the guidelines of the French National Ethics Committee for Sciences and Health report on “Ethical Principles for Animal Experimentation” in agreement with the European Community Directive 86/609/EEC (Apafis #18-185 and #30-959). We used wild-type SWISS mouse pups (C.E Janvier, France) for *ex vivo* experiments. For the *in vivo* experiments, we used mouse pups from the breeding of VGluT2-Cre (a gift from M. Esclapez; (Vong et al., 2011) or Emx1-Cre (a gift from A. de Chevigny; (Gorski et al., 2002) mice with 7- to 8-weeks-old wild-type SWISS females (C.E Janvier, France). All efforts were made to minimize both the suffering and the number of animals used.

### EXPERIMENTAL PROCEDURES AND DATA ACQUISITION

#### Viruses

Anterograde tracing experiments and excitatory opsin expression (Chronos) were achieved using AAV8-Syn-Chronos-tdTomato-WPRE-bGH (pAAV-Syn-Chronos-tdTomato was a gift from Edward Boyden (Addgene viral prep # 62726-AAV8 ; http://n2t.net/addgene:62726 ; RRID:Addgene_62726, (Klapoetke et al., 2014). Inhibitory DREADD expression was induced in a Cre-dependent manner using AAV9-hSyn-DIO-hM4D(Gi)-mCherry (pAAV-hSyn-DIO-hM4D(Gi)-mCherry was a gift from Bryan Roth (Addgene viral prep # 44362-AAV9; http://n2t.net/addgene:44362; RRID:Addgene_44362, (Krashes et al., 2011). Fluorescent protein mCherry expression was induced in a Cre-dependent manner using AAV5-hSyn-DIO-mCherry (pAAV-hSyn-DIO-mCherry was a gift from Bryan Roth (Addgene viral prep # 50459-AAV5 ; http://n2t.net/addgene:50459 ; RRID:Addgene_50459). *In vivo* calcium imaging experiments were performed using AAV1-hSyn-GCaMP6s.WPRE.SV40 (pAAV.Syn.GCaMP6s.WPRE.SV40 was a gift from Douglas Kim & GENIE Project (Addgene viral prep #100843-AAV1; http://n2t.net/addgene:100843; RRID:Addgene_100843, (Chen et al., 2013), AAV9-hSyn-FLEX-axonGCaMP6s (pAAV-hSynapsin1-axon-GCaMP6s was a gift from Lin Tian (Addgene viral prep # 112010-AAV9 ; http://n2t.net/addgene:112010 ; RRID:Addgene_112010, (Broussard et al., 2018) and AAV1-CAG-tdTomato (pAAV-FLEX-tdTomato was a gift from Edward Boyden (Addgene viral prep # 59462-AAV1 ; http://n2t.net/addgene:59462 ; RRID:Addgene_59462)).

#### Virus injections

For P0 stereotaxic virus injections (AAV8-Syn-Chronos-tdTomato-WPRE-bGH, AAV9-hSyn-DIO-hM4D(Gi)-mCherry, AAV5-hSyn-DIO-mCherry or AAV9-hSyn-FLEX-axonGCaMP6s) mouse pups were anesthetized by inducing hypothermia on ice and maintained on a dry ice-cooled stereotaxic adaptor (Stoelting, 51615) fixed to a stereotaxic frame with a digital display console (Kopf, Model 940). Under aseptic conditions, an incision was made in the scalp and the skull was exposed and gently drilled (Ball Mill, Carbide, #¼ .019”, 0.500 mm diameter, CircuitMedic). The ventral midline thalamus (VMT), the entorhinal cortex (EC) and the dorsal hippocampus (dHPC) were targeted by empirically determined coordinates, based on the Atlas of the Developing Mouse Brain (Paxinos et al., 2020). Using the transverse sinus and the superior sagittal sinus as references, VMT coordinates were 1.15 mm anterior from the sinus intersection; 0.1 mm lateral from the sagittal sinus; 2.5 mm depth from the brain surface. The EC injection site was ∼0.62 mm posterior from the sinus intersection; ∼2.73 mm lateral from the sagittal sinus; at 2 mm depth from the brain surface. Finally, dHPC coordinates were 0.8 mm anterior from the sinus intersection; 1.5 mm lateral from the sagittal sinus; 1.1 mm depth from the brain surface. Ten or twenty nL of undiluted viral solution were delivered using a glass pipette pulled from borosilicate glass (3.5” 3-000-203-G/X, Drummond Scientific) and connected to a Nanoject III system (Drummond Scientific). The tip of the pipette was broken to achieve an opening with an internal diameter of 30-40 μm. We injected 10 or 20 nL of AAV8-Syn-Chronos-tdTomato-WPRE-bGH in the VMT or the EC of SWISS mouse pups. Emx1-Cre mouse pups were injected in the EC or the dHPC with 10 nL AAV9-hSyn-DIO-hM4D(Gi)-mCherry. VGluT2-Cre mouse pups were injected in the VMT with 10 nL of either AAV9-hSyn-DIO-hM4D(Gi)-mCherry or AAV9-hSyn-FLEX-axonGCaMP6s.

The intracerebroventricular injection protocol was adapted from published methods (Kim et al., 2014); (Dard et al., 2022). Briefly, mouse pups were anesthetized by hypothermia. Emx1-Cre mouse pups were injected in the left lateral ventricle with 2 µL of a mixture of AAV2.1-hSynGCAMP6s.WPRE.SV40 and Fast Blue dye (1:20). VGluT2-Cre mouse pups were similarly injected with either the AAV2.1-hSynGCAMP6s.WPRE.SV40/Fast Blue dye mixture or a AAV1-CAG-tdTomato/Fast Blue dye (1:20) mixture. To reach the ventricle, we injected in a position that was roughly two fifths of an imaginary line drawn between lambda and the left eye at a depth of 0.4 mm. Correct injection was verified by the spreading of the blue dye.

At least five days were allowed for virus expression before recording procedures.

#### Drugs

Solutions of SR95531 hydrobromide (Tocris, catalogue no. 1262/10) and NBQX disodium salt (Tocris, catalogue no. 1044/10) were made to final concentrations of 10 µM and 20 µM, respectively.

Clozapine N-oxide solution (CNO) was prepared at a final concentration of 1 mg/mL. Briefly, CNO (Tocris, Ref# 4936, 10mg) was dissolved in 5% DMSO and then diluted with saline solution (DPBS, 14190-144, Life Technologies) to reach 1 mg/mL CNO and 0.25% DMSO. Animals were injected once, before *in vivo* recordings, with 10 μL/g of this solution to reach the final dose of 10 mg/kg.

#### Slice Preparation for ex vivo experiments

Horizontal hippocampal slices (400 µm thick) were obtained from 5 to 7-postnatal days old (P5–P7) Chronos-injected and non-injected (control) SWISS pups. Brains were sliced with a Leica VT1200 S vibratome in ice-cold oxygenated modified artificial cerebrospinal fluid containing (in mM): 2.5 KCl, 1.25 NaH_2_PO_4_, 7 MgCl_2_, 5 CaCl_2_, 26 NaHCO_3_, 5 D-glucose, 126 CholineCl. Slices were left for one hour at room temperature in oxygenated ACSF containing (in mM): 126 NaCl, 3.5 KCl, 1.2 NaH_2_PO_4_, 26 NaHCO_3_, 1.3 MgCl2, 2.0 CaCl_2_, and 10 D-glucose.

#### Ex vivo 2-photon calcium imaging and optogenetics

*Ex vivo* 2-photon calcium imaging experiments were performed on injected and non-injected (control) SWISS pups’ slices using the calcium indicator Fura2-AM (Molecular Probes, F1221). Slices were incubated in the dark at 35-37 °C for 30 min within a small petri dish containing 2.5 mL of oxygenated ACSF with 25 µL of a 1 mM Fura2-AM solution (in 96 % DMSO, 4 % pluronic acid). One hour later, slices were transferred to a recording chamber continuously perfused with oxygenated ACSF (3 mL/min) at 35–37°C. Calcium imaging was performed with a multibeam multiphoton pulsed laser scanning system (LaVision Biotech) coupled to a microscope as previously described (Crépel et al., 2007). Images were acquired through a CCD camera, which typically resulted in a time resolution of 124 ms per frame. Slices were imaged using a 20X, NA 0.95 objective (Olympus). Imaging depth was on average 80 μm below the surface (range: 50–100 μm).

Optogenetics stimulation was performed by sending a TTL signal generated with Clampex software and a 1440A Digidata (Molecular Devices) to a 488 µm OBIS laser (Coherent). The light pulses were delivered through an optic fiber placed above the CA1 *stratum lacunosum moleculare (slm)*. Imaging acquisition was divided in 3 equal parts: (1) a baseline period with no light stimulation during which we recorded spontaneous activity of the CA1 region; (2) a stimulation period; (3) a recovery period with no light stimulation. The stimulation protocol consisted of the application of bursts of 3 light pulses (3 ms per light pulse) at 20 Hz. Bursts were repeated at 0.1 or 0.2 Hz.

#### Patch clamp recordings and optogenetics in injected SWISS pups

Patch clamp recordings in injected SWISS pups’ slices were performed using a SliceScope Pro 1000 rig (Scientifica) equipped with a CCD camera (Hamamatsu Orca-05G). Slices were transferred to a recording chamber and continuously perfused with oxygenated ACSF (3 mL/min) at ∼35°C. Patch recording electrodes (4-8 MΩ resistance) were pulled using a PC-10 puller (Narishige) from borosilicate glass capillaries (GC150F-10, Harvard Apparatus) and filled with a filtered voltage clamp intracellular solution containing (in mM): 130 Cs-gluconate, 10 CsCl, 0.1 CaCl_2_, 1 EGTA, 10 HEPES, 2 Mg-ATP, 0.3 Na-GTP (pH 7.3 and ∼280 mOsm), and 0.5 % biocytin. Electrophysiological signals were amplified (Multiclamp 700B), low-pass filtered at 2.9 kHz, digitized at 5-20 kHz and acquired using a Digidata 1440A digitizer and pClamp 10 software (all from Molecular Devices).

An optoLED system (Cairn Research) consisting of two 3.5 WLEDs was used. The 470 nm WLED coupled to a GFP filter cube was employed to stimulate Chronos expressing axons (3 ms-long light pulses). The white WLED coupled to an RFP filter cube was used to visualize the Chronos/tdTomato-expressing axons. Light was delivered using a 40× objective, leading to a light spot size of ∼1 mm, which was able to illuminate the whole hippocampus.

Visually guided patch clamp recordings, in whole-cell configuration, were made from randomly sampled CA1 *slm* putative GABAergic interneurons (*slm*INs) and random CA1 pyramidal cells (PC) in the *stratum pyramidale*. Cell type identity was confirmed *post-hoc* based on morphological features (mainly the shape and location of the dendrites and axons, presence or absence of dendritic spines). To determine the nature of the VMT and EC inputs, one suprathreshold light pulse (0.5 – 1 mW/mm² for *slm*INs, 4 – 8 mW/mm² for PC) was applied to cells clamped at −75 mV and at 0 mV, the reversal potential for GABAergic and glutamatergic transmission, respectively. Since some of the *slm*INs showed monosynaptic GABAergic responses to stimulation of EC afferents, all the recording protocols for *slm*INs with EC stimulation were performed in the presence of the GABA_A_ receptor blocker SR 95531 hydrobromide. To measure the postsynaptic current amplitude and paired pulse ratio on *slm*INs, two pulses of light separated by 50 ms were applied on cells voltage-clamped at −75 mV. To confirm the contribution of an AMPA-R component to the monosynaptic responses at −75 mV, suprathreshold light pulses were applied in the presence of NBQX disodium salt (Tocris, catalogue no. 1044/10). For NMDA/AMPA ratio (N/A ratio) on *slm*INs, the previous stimulation protocol was repeated on the same cell voltage-clamped at +40 mV in the presence of the NBQX disodium salt. To measure the postsynaptic current amplitude of CA1 PCs, one suprathreshold light pulse was applied on cells voltage-clamped at −75 mV. To quantify the polysynaptic responses of CA1 PCs during the stimulation of VMT inputs, recordings were divided in three periods: (1) a baseline period during which we recorded spontaneous postsynaptic currents; (2) a stimulation period; (3) a recovery period with no light stimulation. The one second time period following the first light pulse of the stimulation train (10 pulses total at 20Hz) was considered as the stimulation period. PCs were clamped at −75 mV and at 0 mV. Then, polysynaptic responses on PCs were recorded at −75 mV in the presence of the GABA_A_ receptor blocker, SR 95531 hydrobromide, during the light stimulation protocol of VMT afferents. For all protocols, sweeps were separated by a 20 s delay period to avoid the induction of plasticity and ensure stable responses.

#### Surgery for CA1 in vivo extracellular electrophysiological recordings

Surgery was performed in mice aged between P5 and P8 under isoflurane anesthesia (2-4%). Before surgery, mice received a subcutaneous injection of buprenorphine (0,03mg/kg) and topical administration of lidocaine/prilocaine ointment (5%) on the scalp. During anesthesia, the scalp was removed and the skull cleaned. Two silver chloride wires, serving as ground and reference, were de-insulated at the tip for 1mm, placed between the skull and the dura above the posterior part of the auditory cortex, secured with cyanoacrylate glue (vetbond, 3M) and later fixed with dental cement. A 3 by 1 cm head bar was then fixed above the cerebellum and posterior visual cortex with dental cement, letting the areas above both hippocampi accessible. Craniotomies were then performed at the following coordinates with regards to Bregma: AP = 1,3 mm and ML=1,5 mm between P5 and P6 or AP= 1,5 mm and ML= 1,7 mm between P7 and P8. The dura at the craniotomy sites was then covered with surgical silicone adhesive (kwik-cast) for protection until recording time. Pups were then let to recover without anesthesia on a heating pad.

Thirty minutes after surgery, pups were secured to the recording apparatus by fixing the head bar to the set-up. The rest of the body rested on a heating pad to maintain body temperature at 37°C. CNO at a dose of 10 mg/kg was then injected subcutaneously. Throughout this period and the following experiment, pups were constantly monitored and fed with veterinary milk *ad libitum*.

#### CA1 in vivo extracellular electrophysiological recordings

Before the recordings, the silicone adhesive was removed from the craniotomy and a 32-channel silicon probe (Buzsaki32, Neuronexus), coated with a red fluorescent marker (DiI, Sigma) was lowered in the dorsal CA1 until it reached the pyramidal layer (∼1200 µm depth). The electrophysiological signal from the electrodes was digitized and sampled at 30KHz, filtered (0.1 Hz-6000 Hz) and multiplexed by a preamplifier (RHD-32 channel, Intan Technologies). It was then recorded synchronously with the analog signal from the Piezzo-electric sensors via an acquisition system (RHD2000_evaluation_system, Intan Technologies). Acquisition was controlled via the Open-Ephys (Siegle et al., 2017) graphical user interface. Recordings started 5 minutes after CNO injection and lasted for 1 hour.

#### Surgery for CA1 in vivo 2-photon imaging

The surgery to implant a 3-mm-large cranial window above corpus callosum was performed as already described (Dard et al., 2022) and adapted from previous methods. Briefly, P6-P8 mice were anesthetized and maintained during the whole surgery with 1-3 % isoflurane in a mix of 90% O_2_-10% air. Body temperature was controlled and maintained at 36°C. Analgesia was controlled using buprenorphine (0.05 mg/ kg). After fixing a small custom-made headplate, the skull was removed, and the cortex was gently aspirated until the appearance of the external capsule. At the end of the cortex resection, we sealed a 3 mm-diameter glass circular cover glass (#1 thickness, Warner Instrument) attached to a 3 mm-diameter and 1.2 mm-high cannula (Microgroup INC) using KwikSil adhesive (WPI). The edge of the cannula was fixed to the skull with cyanoacrylate. We let the animal recover on a heated pad for at least one hour before the imaging experiment. Throughout this period and the following experiment, pups were constantly monitored and fed with veterinary milk *ad libitum*.

#### CA1 in vivo 2-photon calcium imaging in mouse pups

Two sets of *in vivo* imaging experiments were performed: (1) calcium imaging of VMT axons in the CA1 *slm* area (VGluT2-Cre mouse pups injected with AAV9-hSyn-FLEX-axonGCaMP6s in VMT and AAV1-CAG-tdTomato in the left lateral ventricle), and (2) calcium imaging of the CA1 neurons in *the stratum pyramidale* with DREADD inhibition of VMT or EC (VGluT2-Cre or Emx1-Cre mouse pups injected with AAV9-hSyn-DIO-hM4D(Gi)-mCherry, respectively, in the VMT or the EC, and AAV1-hSynGCaMP6s.WPRE.SV40 in the left lateral ventricle). Two-photon calcium imaging was performed on the day of the window implant using a single beam multiphoton pulsed laser scanning system (TriM Scope II, LaVision Biotech) coupled to a microscope as previously described (Villette et al., 2015). Images were acquired through a GaAsP PMT (H7422-40, Hamamatsu) using a 16X water immersion objective (NIKON, NA 0.8).

For calcium imaging of VMT axons in CA1 *slm* region (1), imaging fields were located 300 to 400 µm below the *stratum pyramidale*, i.e. 700 µm under the glass window. Using Imspector software (LaVision Biotech), the fluorescence signal from a 400 μm x 400 µm field of view (FOV) was acquired at approximately 2 Hz with a 6.7 μs dwell time per pixel (2 μm/pixel) or from a 300 µm x 300 µm FOV at approximately 1 Hz with a 0.4 μs dwell time per pixel (0.29 μm/pixel)

For DREADD experiments (2), FOVs were selected to sample the intermediate CA1 area and maximize the number of imaged neurons in the *stratum pyramidale*. FOVs of 400 μm x 400 µm were acquired at approximately 8 Hz with a 1.8 μs dwell time per pixel (2 μm/pixel). Ten-minute imaging blocks were performed for one hour following the subcutaneous injection of CNO, at a dose of 10 mg/kg, or saline solution (DPBS, 14190-144, Life Technologies).

Simultaneously with imaging experiments, mouse pup motor behavior was monitored using two infrared cameras (Basler, acA1920-155um) positioned on each side of the animal. For each camera, a square signal corresponding to the exposure time of each frame from the camera was acquired and digitized using a 1440A Digidata and the Axoscope 10 software (Molecular Devices).

The success rate of this experiment depends on the combined success of the P0 stereotaxic virus injections and the window implant surgery. We performed this on 2 mice out 18 for the VMT axons imaging experiments in the CA1 *slm*region and 11 mice out of 64 (EC: 5/36; VMT: 6/28) for the DREADD experiments.

#### Histological processing

For anatomical validation of the stereotaxic injections in VMT and EC, 50 µm brain slices from P7 mice injected either in EC or in VMT at P0 were processed as described previously (Dard et al., 2022). Briefly, under deep anesthesia (Domitor and Zoletil, 0.9 and 60 mg/kg, respectively) pups were transcardially perfused with 4% paraformaldehyde (PFA) in 0.1M PBS (PBS tablets, 18912-014, Life technologies). Brains were post-fixed overnight at 4°C in 4% PFA in 0.1M PBS, washed in PBS, cryo-protected in 30% sucrose in PBS, before liquid nitrogen freezing. Brain slices were obtained by using a cryostat (CM 3050S, Leica) and collected on slides. Then, they were processed for immunocytochemical labelling. Briefly, sections were bathed in PBS-0.3% Triton (PBST) and 10% normal donkey serum (NDS), then incubated overnight with the primary antibody rabbit anti-dsRed (1:1000 in PBST; Clontech, AB_10013483) at 4°C. After several washes with PBST, slices were incubated 2 hours at room temperature with a secondary antibody donkey anti-rabbit Alexa 555 (1:500 in PBST, ThermoFisher, A31570). After Hoechst counterstaining, slices were mounted in Fluoromount and kept at 4°C until microscope acquisition.

For the Emx1-Cre and VGluT2-Cre mice used for the *in vivo* experiments, brains were directly post-fixed overnight at 4 °C in 4% PFA in 0.1M PBS, washed in PBS, sectioned using a vibratome (VT 1200 s, Leica) into coronal 150 μm-thick slices and collected on slides. Then immunocytochemistry was performed as previously described to amplify mCherry signal.

For biocytin-filled patched cells, slices were fixed overnight at 4 °C in 4% PFA in 0.1M PBS, rinsed in PBS, cryo-protected in 30% sucrose in PBS, before freezing within liquid isopentane cooled with dry ice. Slices were incubated with 10% NDS in PBST and then in a mix containing streptavidin-488 (1:2000 in PBST) and primary antibodies rabbit anti-dsRed (1:2000 in PBST) for 5 days at room temperature. As described above, slices were then incubated 4 hours at room temperature with secondary antibody donkey anti-rabbit Alexa 555 (1:500 in PBST) and mounted in Vectashield® with DAPI (H-1200, Vector).

For brain tissue clearing, fixed brains from P7 mice injected with AAV8-Syn-Chronos-tdTomato-WPRE-bGH either in EC or in VMT at P0 were used. We used Animal Tissue-Clearing Reagent CUBIC kit (CUBIC-L [T3740] and CUBIC-R [T3741], Tokyo Chemical Industry). Briefly, fixed brains were incubated in a CUBIC-L solution diluted by half in distilled water (50 % CUBIC-L) at room temperature overnight. Then brains’ lipids were removed by immersing them in pure CUBIC-L solution for 5 days at 37°C. The solution was changed every 2 days. After 5 days, brains were washed in 0.1 M PBS solution for a whole day. Then, brains were incubated in a CUBIC-R solution diluted by half in distilled water (50 % CUBIC-R) at room temperature overnight and in the pure CUBIC-R solution until the acquisition within a 24h delay.

#### Image acquisition

Epifluorescence images were obtained with a Zeiss AxioImager Z2 microscope coupled to a camera (Zeiss AxioCam MR3) with an HBO lamp associated with 470/40, 525/50 and 545/25 filter cubes. Confocal images were acquired with a Zeiss LSM-800 system equipped with a tunable laser providing an excitation range from 405 to 670 nm. For the analysis of the proximo-distal and radial distribution of VMT and EC inputs in CA1 region, 5 µm thick stacks were taken from the surface (z=1 µm, pixel size=0.3 µm) with the confocal microscope using a Plan-Apochromat 20x/0.8 objective. For the analysis of EC inputs distribution in the CA1 region of patched-cell slices, tiles were acquired with the epifluorescence microscope using the ApoTome.2 module (Zeiss) focused on the patched - cell soma using an EC Plan-Neofluar 10x/0.30 M27 objective. For the analysis of the number of contacts made by labelled thalamic or entorhinal afferents on biocytin filled CA1 *slm* GABAergic cells, confocal stacks centered on the soma and neurites of biocytin filled cells were acquired at constant resolution (0.065 μm/pixel) and z-step (0.5 μm) with a plan-achromat 40x/1.4 oil using a 2,4x scan zoom.

Whole cleared brain acquisitions were performed using the UltraMicroscope II (LaVision Biotec) coupled with a superK EXTREME (NKT photonics) Supercontinuum laser. Excitation wavelength was selected with band pass filters (470/40 and 560/40) and converted as a light sheet of 4 µm thickness. Two horizontal z-stacks, one for each hemisphere, were taken with a 2x objective (NA: 0.5) plus a x0.63 optical zoom (spatial resolution: 4.52×4.52×4.6 µm). Z-stacks were merged using the pairwise stitching plugin on Fiji.

### DATA PRE-PROCESSING

#### Ex vivo 2-photon calcium imaging pre-processing

We used custom designed MATLAB software allowing for the segmentation and extraction of calcium traces from labelled cells. Because, for each light stimulation, 1 or 2 frames were saturated, we replaced the saturated frames by the 1 or 2 frames that preceded them. Then, onsets and offsets of calcium signals were detected using another custom designed MATLAB program (Bonifazi et al., 2009). Network synchronizations (GDPs) were detected as synchronous onsets peaks including more neurons than expected by chance, as previously described (Bonifazi et al., 2009).

#### In vivo extracellular electrophysiological recordings pre-processing

Multi-unit activity (MUA) was detected from bandpass-filtered (300-6000 Hz; two successive zero-phase, 3rd and 5th order Chebychev filters, respectively) local field potentials (LFP) recorded from each channel. MUA action potentials were detected when the signal reached a threshold equal to 4,5 times the estimated standard deviation from the signal distribution. Any activity that was detected simultaneously on more than four channels was considered as an artifact and discarded. MUA frequency across time was then estimated in 0.5 s bins; smoothed using a 30 s moving average filter and normalized to the mean MUA frequency measured in the first 10 minutes of recording.

#### In vivo 2-photon calcium imaging data pre-processing

##### Motion correction

Image series were motion corrected using the NoRMCorre algorithm available in the CaImAn toolbox (Pnevmatikakis and Giovannucci, 2017).

##### Cell segmentation

Cell segmentation was achieved using Suite2p (Pachitariu et al., 2017). Axonal branch segmentation was performed using Suite2p or Fiji (http://fiji.sc) where ROIs were manually defined.

##### Cell type prediction

Cell type prediction was done using the DeepCINAC cell type classifier (Denis et al., 2020). Briefly, a neuronal network composed by CNN (Convolutional Neural Network) and LSTM (Long Short Term Memory) was trained using labelled GABAergic neurons, pyramidal cells and noisy cells to predict the cell type using 100 frames long movie patches centered on the cell of interest. Each cell was classified as GABAergic neuron, pyramidal cell or noise. Cells classified as ‘noisy cells’ were removed for further analysis.

##### Activity inference

For 2-photon calcium imaging of the pyramidal cell layer, activity inference was done using DeepCINAC classifiers (Denis et al., 2020). Briefly, a classifier composed by CNN and LSTM was trained using manually labelled movie patches to predict neuronal activation based on movie visualization. Depending on the inferred cell type, activity inference was done using either a general classifier or an interneuron specific activity classifier. Activity inference results in a “cells x frames matrix” giving the probability for a cell to be active at any single frame. We used a 0.5 threshold on this probability matrix to obtain a binary activity matrix considering a neuron as active from the onset to the peak of a calcium transient. For 2-photon calcium imaging of VMT axonal branches in the CA1 *slm*, ΔF/F was calculated and a Gaussian low-pass filter was applied to denoise the trace. Frames with z-movement were excluded from the traces to determine the calcium transients’ threshold. This threshold corresponded to the median of the free z-movements denoised trace plus 3 times its interquartile range. Significant transients were conserved if their onset was out of a z-movement period.

##### Behavior analysis

Analysis of video tracking was done using CICADA and behavior was manually annotated in the BADASS (Behavioral Analysis Data And Some Surprises) GUI (Denis et al., 2020). Based on the video tracking three categories of movements were described: complex movements, startles and twitches (irrespectively of the limb).

##### Neurodata without border (NWB) embedding

To be analyzed, all of the preprocessed data were combined into a single NWB:N (Rübel et al., 2022) file per imaging session containing imaging data and metadata, cell contours, cell type prediction, neuronal activity inference and animal behavior.

### DATA ANALYSIS

#### Ex vivo 2-photon calcium imaging and optogenetics analysis

The analysis of the photostimulation effect on network activity is based on the computing of Inter GDP Intervals (IGI) and stimulation-associated-GDPs. To establish whether the photostimulation of external inputs was able to influence the frequency of GDPs, we calculated the IGI during the photostimulation period and the non-stimulation periods (control). To assess whether a phasic light stimulation of the VMT or EC inputs was able to lock GDP occurrence within one second following the stimulation (same time window as previously used in (Bonifazi et al., 2009)), we counted the number of GDPs occuring within a 1 second time window after light stimulations. To evaluate whether the number of GDPs following light stimulations was higher than by chance, we compared this value to the 99th percentile of the distribution of the number of GDP following light simulation obtained from 1000 surrogates.

#### Synchronous Calcium Event (SCE) detection in *in vivo* recordings

Synchronous Calcium Events were defined as the imaging frames in which the number of co-active cells exceeded the chance level as estimated using a reshuffling method as described previously (Dard et al., 2022). The SCE detection threshold, which corresponds to the minimal number of significantly co-active cells, was determined using the first ten minutes of the recording and applied to the other ten-minute imaging blocks.

#### Patch clamp recordings and optogenetics analysis of ex vivo recordings

The analysis of the monosynaptic responses evoked by the light stimulation of EC or VMT inputs on CA1 *slm* GABAergic neurons and pyramidal cells was performed using Clampfit 10 (Molecular Devices). We measured the delay between the onsets of the light pulse and the cell response and its amplitude. The paired pulse ratio (PPR) and the NMDA/AMPA ratio (N/A ratio) was calculated based on the amplitude measures. For the analysis of the polysynaptic responses evoked by the light stimulation train applied to external inputs, we used the TaroTools toolbox (https://sites.google.com/site/tarotoolsregister) for IgorPro (WaveMetric). This allowed us to extract the time stamps of the PostSynaptic Currents (PSCs). Then, the polysynaptic effect of the stimulation train was estimated with the Peri-Stimulus Time Histogram (PSTH) performed with a MATLAB custom-made script.

#### Distribution of VMT and EC afferents in the CA1 region

Analysis of the anatomical distribution of VMT and EC putative axons in the CA1 area was performed on confocal or epifluorescence images using Fiji (http://fiji.sc). First, tissue auto-fluorescence was removed by subtracting the mean value of a square ROI placed on the CA3 region. For axonal density measurements along the radial axis, a line was drawn from the alveus to the hippocampal fissure in the middle of the CA1 region and the plot profile plugin was applied. For the axonal density measurement along the proximo-distal axis, the CA1 *slm* ROI was defined manually, straightened out and the plot profile was done along the proximo-distal axis. For each profile plot, the distance was given as a percentage relative to the maximal distance; the intensity profile was normalized to the peak.

#### EC and VMT putative contacts onto CA1 slm neurons

Appositions between thalamic or entorhinal tdTomato positive axons and biocytin-filled somata and dendrites, considered as putative contacts, were counted manually throughout z-stacks using the multi-point tool plugin in Fiji (http://fiji.sc). The total red and green surface per cell was calculated as the sum of the red and green surfaces measured for each stack using the isodata threshold Fiji plugin.

#### Statistics

Statistical tests were performed using GraphPad (Prism). We used either two-tailed Mann-Whitney, Student t-test or Welch t-test, depending on the results for normality and homoscedasticity tests, to compare data from axonal density experiments, *ex vivo* calcium imaging and patch-clamp recordings. For *in vivo* 2-photon calcium imaging, cell transients frequency was compared between the 0-10 minutes and the 40-50 minutes using Wilcoxon test. The number of SCEs was compared between the two periods of the 3 experimental conditions (EC, VMT and Ctrl) using two-way ANOVA.

## RESULTS

### Spatial distribution of VMT and EC inputs onto the CA1 region at early postnatal stages

The distribution of extra-hippocampal axons in the CA1 region has been described in adult mice. In particular, it is well established that VMT inputs target mainly the ventro-intermediate part of CA1 (Hoover and Vertes, 2012). In order to determine the possible influence of VMT and EC inputs onto early CA1 dynamics, we first analyzed their distribution along the radial (in the CA1, from *alveus* to *slm*) and proximo-distal (in CA1 *slm*, from CA2 to subiculum) anatomical axes, in the ventro-intermediate CA1 region. To this aim, mouse pups were stereotactically injected with a virus expressing tdTomato (together with the fast opsin Chronos (Klapoetke et al., 2014); (Ronzitti et al., 2017) for later optogenetic experiments) either in the ventral midline thalamus (VMT, **Figure S1**) or in the entorhinal cortex (EC, **Figures S2A, S2B**). Injections were performed at birth (P0, **Figure 1A**). We analyzed the pattern of innervation of CA1 by VMT and EC in fixed brain slices (five to seven days after injection) and found that both inputs innervated the *slm* (**Figure 1B**, **Supplementary Movies 1 and 2**). As expected (Supèr and Soriano, 1994), EC axons were also visible in the *alveus* as well as in the contralateral CA1 *slm* (**Figure S2C**, **Supplementary Movie 1**); VMT projections are mainly present in the ventro-intermediate CA1 (**Supplementary Movie 2**). These observations are consistent with previous studies (Hartung et al., 2016; Supèr and Soriano, 1994). EC inputs were present all along the proximo-distal axis in the CA1 *slm*, but innervated more densely the proximal part of the CA1 *slm*, at the CA2 border (n=4 mice, 18 slices; **Figure 1C, left panel**). Conversely, VMT inputs innervated the distal CA1 *slm*, at the border with the subiculum (n=4 mice, 16 slices; **Figure 1C, left panel**). The spatial segregation of EC and VMT inputs along the proximo-distal axis was quantified by measuring the relative distance of the peak of axonal fluorescence from the distal limit of CA1 (border with the subiculum) (VMT: mean=20 ± 8 %, n=16 slices, 4 animals, EC: mean=69 ± 19 %, n=18 slices, N=4 animals, Welch’s t-test, p<0.0001; **Figure 1C, right panel**). On the radial axis, EC and VMT inputs were mostly concentrated in the CA1 *slm* (**Figures 1B and D**). This anatomical segregation of EC and VMT inputs suggests that, if active, they may influence different CA1 circuits during early postnatal development.

**Figure 1.**
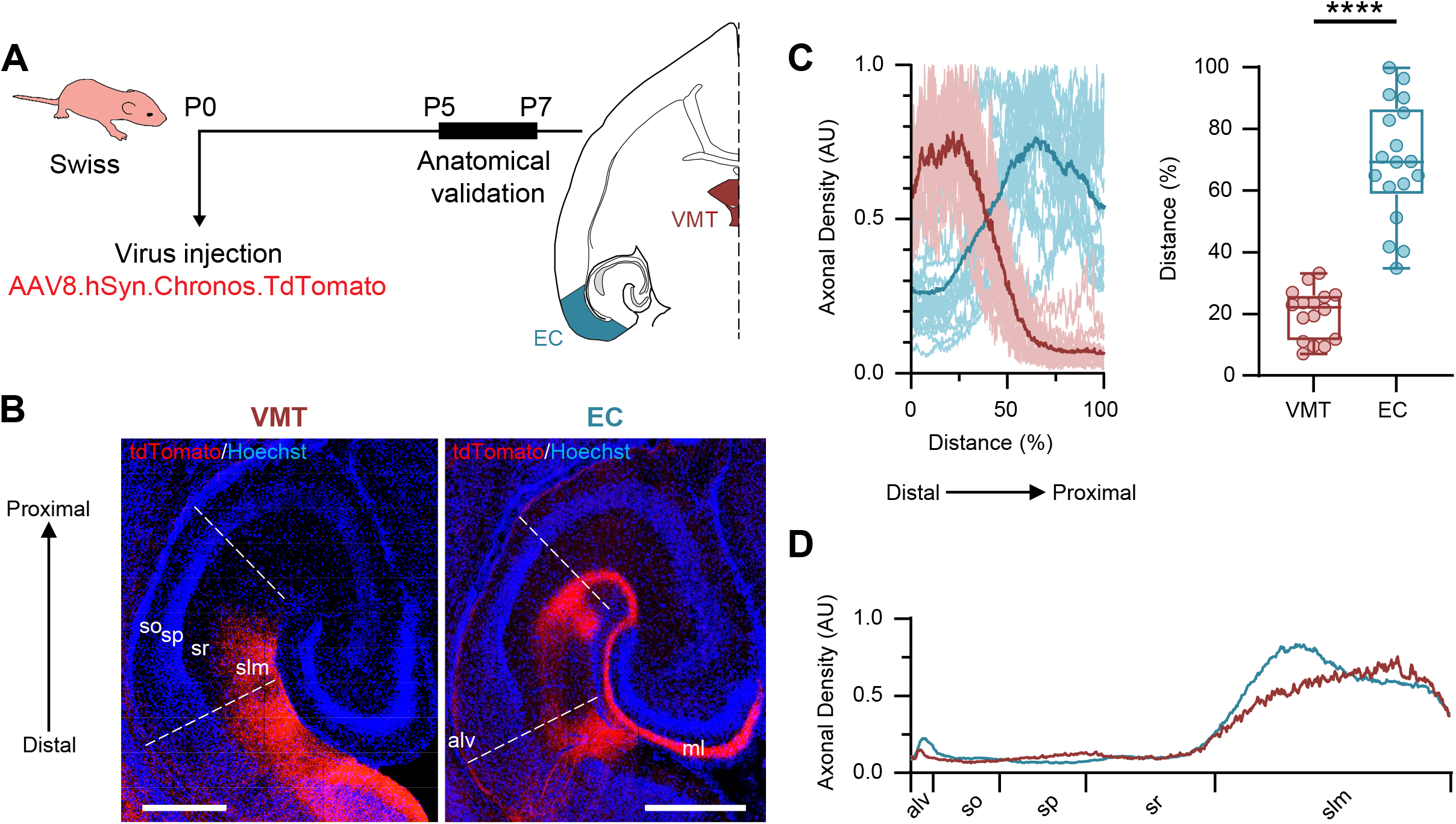
Anatomical segregation of thalamic and entorhinal inputs in the CA1 *slm* during development. **(A)** Schematic representation of the experimental timeline. **(B)** Confocal images of horizontal sections of the ventral part of P7 mice hippocampus showing the distribution of labeled fibers after tdTomato virus injection in the VMT (left) or EC (right). DAPI staining was used to define the different layers of CA1 region: *alveus* (*alv*), *stratum oriens* (*so*), *stratum pyramidale* (*sp*), *stratum radiatum* (*sr*) and the *stratum lacunosum moleculare* (*slm*). The dashed lines indicate the distal border, close to the subiculum, and the proximal border, close to CA2. VMT fibers were present in the CA1 *slm* while EC afferents were observed in the alveus, the CA1 *slm* and the molecular layer (ml) of the dentate gyrus. Scale bar: 400 µm. **(C)** Left: Distribution of VMT (red) and EC (blue) axon density in CA1 *slm* along the disto-proximal axis (0 to 100 %). Each light coloured curve represents the normalized measurement from an individual slice (VMT (red) : n=16 slices, N=4 animals, EC (blue) : n=18 slices, N=4 animals). Dark coloured curves correspond to the mean of all light curves. Right: Distance from the distal border of VMT and EC axonal density peaks. Each dot corresponds to the location from the distal border of the axonal intensity peak for a slice. Each boxplot shows the 25th, 50th and 75th percentiles and whiskers represent the 5th to the 95th percentile. Significant difference between VTM (n=16 slices, N=4 animals) and EC (n=18 slices, N=4 animals) was observed (Welch’s t-test, t=9.928, p<0.0001). **(D)** Mean distribution of VMT (red, n=16 slices, N=4 animals) and EC (blue, n=17 slices, N=4 animals) axon density in the CA1 radial axis.

### Modulation of local CA1 dynamics by photostimulation of EC or VMT inputs

While a recent study has shown that the EC is active in neonatal rodents (Valeeva et al., 2019), it remains unclear whether the VMT input is also active at this developmental stage. To address this question before the stimulation experiments, we performed *in vivo* 2-photon calcium imaging from VGluT2-expressing axons in the *slm* in neonatal mouse pups. We used virus injections at birth to express the axon-enriched calcium indicator protein GCaMP6s (Broussard et al., 2018) Cre-dependently in VGluT2-Cre mice, and td-Tomato (independently from Cre) to locate the *slm* (see methods; **Figure S3A-C**). Since VGluT2 tags subcortical excitatory neurons (including thalamic neurons), calcium activity in VGluT2+ fibers in the *slm* largely reflects activity of the VMT input. We observed spontaneously occurring calcium transients about once every 5 minutes in boutons expressing GCaMP6s (median=0.24 transients/minute [iqr=0.34]; **Figures S3D, S3E**). We conclude that together with activity originating from EC, VMT inputs can impact CA1 dynamics during early postnatal development.

In order to test the influence of these extrinsic inputs onto local CA1 dynamics, we stimulated VMT or EC axonal fibers in hippocampal slices using an optogenetic approach. We focused on the modulation of the Giant Depolarizing Potentials (GDPs), the main type of network activity pattern spontaneously occurring in hippocampal slices between P5 and P7 (Ben-Ari et al., 1989). This pattern can be considered as “internally-generated” as it is embedded in local circuits, being observed in the disconnected CA1 *ex vivo*. We performed calcium imaging combined with CA1 *slm* photo-activation of external afferents in horizontal slices from P5-7 SWISS mice. The slices were loaded with the calcium indicator Fura2-AM as previously described (Crépel et al., 2007). Pups were injected at birth with a Chronos/tdTomato virus in the EC or VMT, as described above. An optic fiber placed above the CA1 *slm* was used for photoactivation of either pathway using blue light. Neuronal activity was recorded in the CA1 pyramidal cell layer using 2-photon calcium imaging (**Figure 2A, Supplementary Movies 3 and 4**). As expected spontaneous synchronous activation of a large portion of imaged cells corresponding to GDPs could be observed ((Crépel et al., 2007) **Figures 2B1, 2C1**). Photoactivation of VMT axons affected the time interval between consecutive GDPs (Inter GDP Interval, IGI) in 5 out of 8 slices (n=7 animals; **Figures 2B2, 2D1**). In these 5 slices, GDPs occurred at a higher frequency since a decrease in IGI during photoactivation was observed in 4 slices and an increase in one (**Supplementary Tables 1 and 2**). Similarly, photoactivation of EC terminals affected IGIs in 4 out of 7 slices, with a decrease in IGI observed in three cases but one (n=5 animals; **Figures 2C2, 2D1, Supplementary Tables 1 and 2**). We conclude that the photostimulation of VMT or EC inputs modulates the occurrence of CA1 GDPs, with an increase in GDP frequency being more frequently observed.

**Figure 2.**
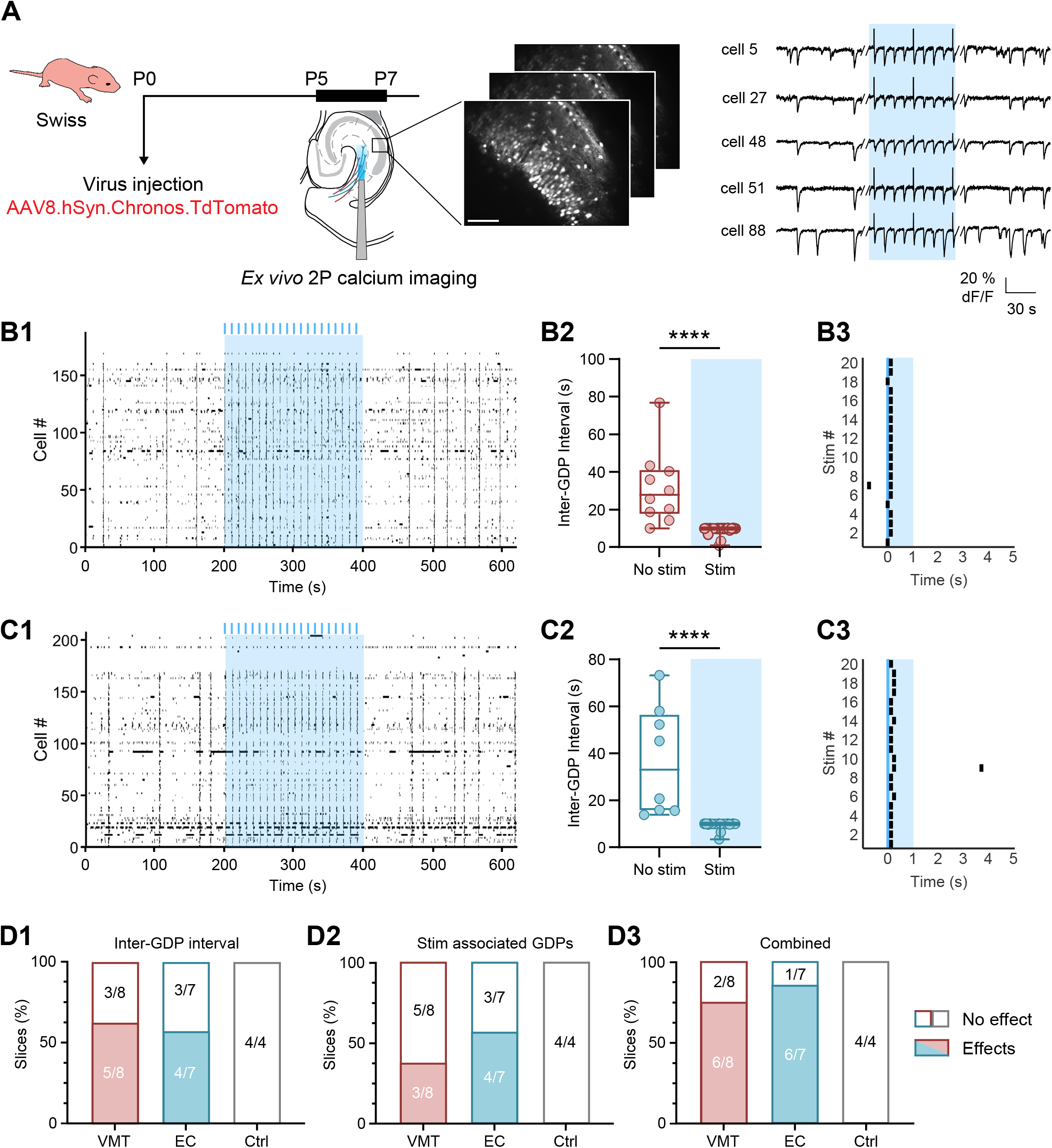
VMT and EC inputs modulate *ex vivo* network activities in the developing CA1. **(A)** Schematic representation of the experimental timeline. On the right, calcium transients (negative variation of dF/F in Fura2-AM) were observed during periods of non-stimulation and during the application of a stimulation train at 0.1 Hz or 0.2 Hz (blue). Scale bar: 50 µm. **(B)**. Example of the effect of VMT afferences photostimulation on the CA1 network activity observed in a P5-P7 brain slice of a wild-type SWISS mouse pup, injected with Chronos/tdTomato in the VMT at P0. B1, Raster plots showing neuronal activity as a function of time. The light blue rectangle corresponds to the photostimulation period during which phasic stimulations at 0.1 or 0.2 Hz were applied (blue lines). B2, Measurement of the inter-GDP interval during the period of stimulation of VMT afferents (light blue) and non-stimulation corresponding to the raster plot shown in 1. Each dot corresponds to the measured IGIs during non-stimulation (“No-Stim”) and stimulation (“Stim”) periods. Each boxplot shows the 25th, 50th and 75th percentiles and whiskers represent the 5th to the 95th percentile. Significant difference between non-stimulation and stimulation periods was observed (No-Stim : median=27.84 s [iqr=66.71], n=10 IGIs, Stim : median=9.920 s [iqr=9.30], n=20 IGIs, Mann-Whitney, U=3.0, p<0.0001). B3, GDPs occurence time with respect to the photostimulation during the stimulation period that correspond to the raster plot shown in 1. GDP occurrence (black rectangles) as a function of time following 0.1 Hz phasic stimulations of VMT afferents. Twenty consecutive light stimulations were centered to time 0 (blue line). The 20 stimulations were followed by GDPs in the following second (light blue rectangle, 99^th^ percentile=17). **(C)** Same as **(B)** but for Chronos/tdTomato injection in P0 wild-type SWISS mouse pups’ entorhinal cortex. C1, Raster plots showing neuronal activity as a function of time. C2, For IGI, significant difference between non-stimulation and stimulation periods was observed (No-Stim : median=32.98 s [iqr=59.27], n=8 IGIs, Stim : median=9.920 s [iqr=6.696], n=20 IGIs, Mann-Whitney, U=0.0, p<0.0001). C3, The 20 stimulations triggered the GDPs in the following second (light blue rectangle, 99^th^ percentile=16.5). **(D)** Proportions of VMT, EC and Control (Ctrl; laser stimulation on slices without virus, see Figure S5C) displaying significant effects on GDP frequency and GDP synchronization upon stimulation. “Combined” corresponds to traces that show an effect on frequency and/or timing of the GDPs.

We next asked whether such photoactivation could directly trigger synchronization of neuronal activity by examining whether GDPs had a higher probability than chance to occur in the one second window following light stimulation. We observed that GDPs were paced by VMT in 3 out of 8 slices (**Figures 2B3, 2D2, S4A**) and in 4 out of 7 slices in the case of EC inputs photoactivation (**Figures 2C3, 2D2, S4B**). Of note, CA1 dynamics were not affected by light in slices that did not express the opsin (n=4 slices, N=2 animals, **Figure S4C**). Detailed results are reported in Supplementary Table 1 and 2. Together, this shows that VMT and EC inputs can modulate spontaneous “internally-generated” CA1 network activity (**Figure 2D3**). We next dissected the local CA1 circuits recruited by VMT and EC inputs.

### VMT and EC inputs recruit different CA1 local circuits

We first examined how EC and VMT terminals impinge onto CA1 *slm* GABAergic neurons. To this aim, we performed patch-clamp recordings from CA1 *slm* neurons in hippocampal slices from P5-7 mice injected with the Chronos/tdTomato virus in the EC or VMT at birth as above (**Figure 3A**). We first examined the nature of VMT and EC synaptic transmission. At the reversal potential for excitatory postsynaptic currents (EPSCs), photostimulation of EC but not VMT inputs could evoke monosynaptic GABAergic responses in some CA1 *slm* GABAergic neurons (7/29 cells; **Figure S5A**), indicating a small but significant contribution of GABAergic transmission to long-range communication from EC as early as P5-7, as in the adult (Basu et al., 2016; Melzer et al., 2012). Thus, we recorded EPSCs evoked by stimulation of the EC inputs in the presence of SR95531, a GABA_A_R blocker, in order to measure the impact of glutamatergic transmission-mediated inputs. Recorded cells distributed along the disto-proximal axis of CA1 *slm* (**Figure 3B**) in locations with detectable tdTomato signal (**Figures S5C, S5D left panels**). Photostimulation of both VMT and EC inputs evoked short-delay EPSCs (monosynaptic glutamatergic responses) in more than two-thirds of the CA1 *slm* neurons (VMT: 68 %, n=47 cells, EC: 67 %, n=34 cells; Delay: VMT: 3.9 ms, n=11 cells, EC: 3.3 ms, n=10 cells; **Figures 3C, S5B left panel**). Moreover, the cells responding to the photostimulation of either afferents were more densely decorated with putative contacts than non-responding cells (VMT: With EPSCs: median=0.18 contacts/stack [iqr=0.13], n=14 cells, Without EPSC: median=0.09 contacts/stack [iqr=0.23], n=11 cells, Mann-Whitney test, p=0.0014; **Figure S5C, middle panel**) or EC (With EPSCs: median=0.15 contacts/stack [iqr=0.28], n=17 cells, Without EPSC: median=0.05 contacts/stack [iqr=0.07], n=9 cells, Mann-Whitney test, p=0.0035; **Figure S5D, middle panel**). Such difference was not due to different background signal as for both inputs, the ratio between cell (green) and axon volume (red) did not significantly differ between responding- and non-responding cells (VMT: With EPSCs: median=0.19 [iqr=0.64], n=14 cells, Without EPSC: median=0.14 [iqr=0.19], n=11 cells, Mann-Whitney test, p=0.0954, **Figure S5C, right panel**; EC: With EPSCs: median=0.11 [iqr=3.59], n=16 cells, Without EPSC: median=0.72 [iqr=4.53], n=9 cells, Mann-Whitney test, p=0.3014, **Figure S5D, right panel**). Morphological inspection of the recorded neurons that were filled with neurobiotin indicated that these cells qualified as putative GABAergic interneurons. We could distinguish putative Neurogliaform cells (NGF) from other CA1 *slm* GABAergic neurons (*slm*INs; **Figure 3D**), based on anatomical criteria, which included their dense axonal arborization covering a larger territory than their short dendrites (Chittajallu et al., 2017; Tricoire et al., 2010). The fraction of *slm* interneurons responding to VMT or EC stimulation was comparable (Fisher’s exact test, p>0.9999, **Figure 3C**). Among these *slm* interneurons, NGF cells were the most likely to respond (NGF: 86 %, n=28 cells, *slm*INs: 51%, n=37 cells ; Fisher’s exact test, p=0.0042, **Figure S5E**). We conclude that CA1 *slm* interneurons, and notably the NGF cells, have an equal probability to be targeted by EC or VMT inputs. These responses to both inputs were mediated by AMPA- and NMDA-R activation (**Figure 3E**) but with EC afferents photostimulation evoking AMPA-R-mediated PSCs of higher amplitude than VMT (VMT: median=24 pA [iqr=121.2], n=10 cells, EC: median=71 pA [iqr=903.1], n=11 cells, Mann-Whitney test, p=0.0610; **Figure 3F**). Two groups of cells could be distinguished based on the amplitude of their AMPA-R-mediated responses to EC afferent photostimulation, one displaying small amplitude EPSCs (median=28 pA [iqr=58.0], n=6 cells) and the other large amplitude EPSCs (median=719 pA [iqr=386.5], n=5 cells, Mann-Whitney test, p=0.0043; **Figures 3E, 3F**). The cells displaying large EPSCs were closer to the hippocampal fissure and CA2 than the small EPSC ones (Radial axis: small EPSCs: median=0.68 [iqr=0.16], n=6 cells, large EPSCs: median=0.79 [iqr=0.09], n=5 cells, Mann-Whitney test, p=0.0173, **Figures 3G, 3H**; Disto-proximal axis: small EPSCs: median=0.48 [iqr=0.25], n=6 cells, large EPSCs: median=0.85 [iqr=0.22], n=5 cells, Mann-Whitney test, p=0.0043, **Figures 3G, 3I**). The presynaptic release probabilities of both afferents were comparable (EC: median=1.02 [iqr=2.19], n=11 cells, VMT: median=0.81 [iqr=2.75], n=10 cells, Mann-Whitney test, p=0.55; **Figure S5B, middle panel**). Similarly, the ratios between NMDA- and AMPA-R mediated PSCs were similar (EC: median=2.05 [iqr=11.81], n=9 cells, VMT: median=2.14 [iqr=3.90], n=9 cells, Mann-Whitney test, p=0.9314; **Figure S5B right panel**). Altogether, these results indicate that both EC and VMT inputs project directly onto CA1 *slm* interneurons, including NGF cells as early as the end of the first postnatal week, with EC inputs innervating preferentially cells located in the proximal part of the CA1 *slm*.

**Figure 3.**
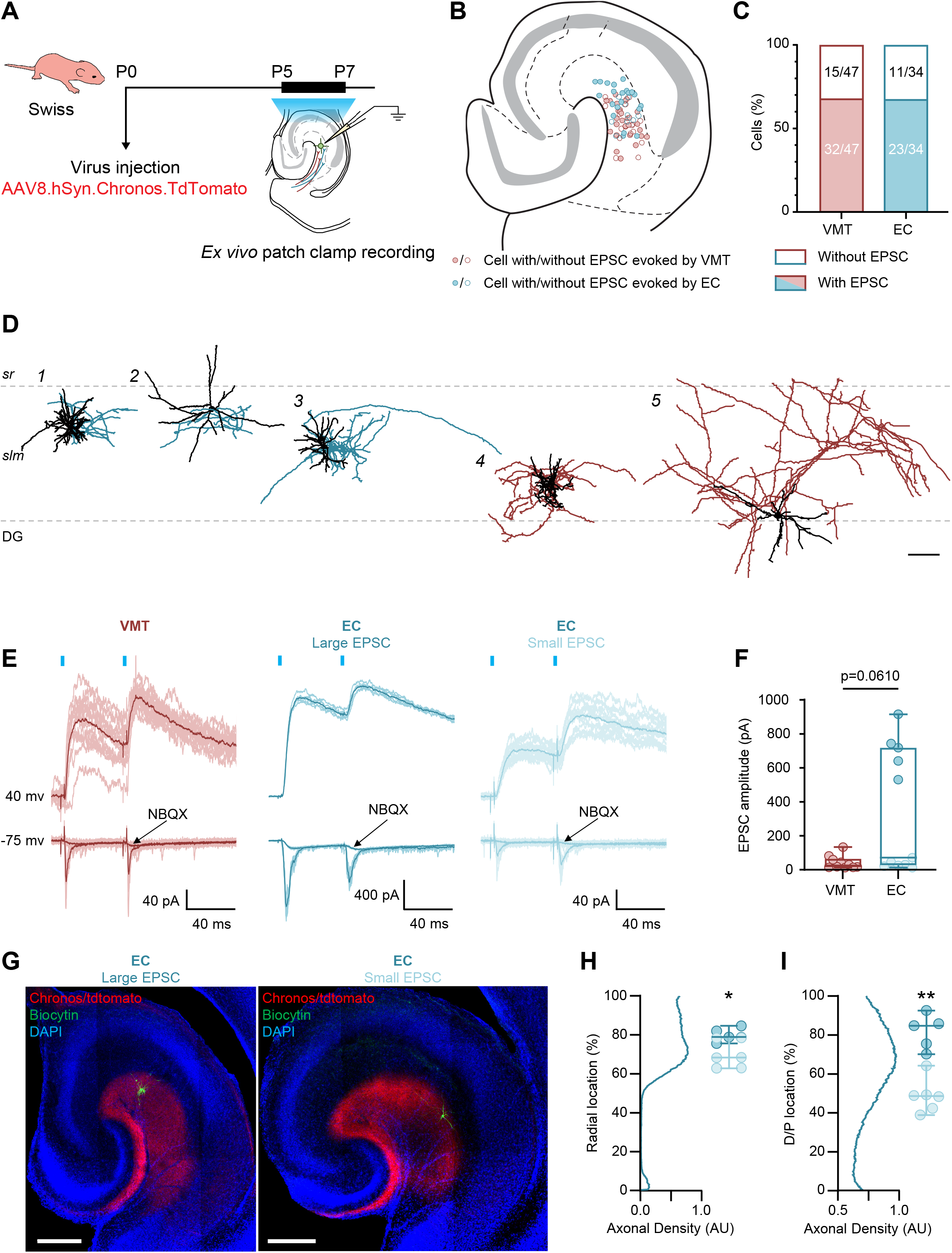
Recruitment of CA1 *slm* GABAergic neurons by VMT and EC inputs during the first postnatal week. **(A)** Schematic representation of the experimental timeline. **(B)** Position of patched CA1 *slm* GABAergic neurons on a schematic representation of the horizontal section of the ventral hippocampus (Adapted from (Franklin and Paxinos, 2007). Filled dots correspond to responding cells to VMT (red) or EC (bleu) photostimulation. Empty dots correspond to no responding cells. **(C)** Proportion of CA1 *slm* GABAergic cells displaying EPSCs upon stimulation of VMT (red, 68.09 %) and EC (blue, 67.65 %) afferents. Empty parts represent cells that didn’t respond to the stimulation (VMT: 31.91 %, EC: 32.35 %). No significant difference between VTM (n=47 cells) and EC (n=34 cells) was observed. **(D)** Neurolucida reconstructions of representative neurobiotin-filled *slm* GABAergic neurons. Axon is depicted in blue for EC-stimulated cells and in red for VMT-stimulated cells. Soma and dendrites are colored in black. Cells 1, 3 and 4 had a neurogliaform-like morphology (NGF) and cells 2 and 5 presented a no-neurogliaform-like morphology (*slm*INs). Scale bar: 100 µm. **(E)** From left to right, monosynaptic glutamatergic responses evoked by the VMT (red) and the EC (blue) LED photostimulation (two light pulses of 3 ms, blue lines). EC photostimulation evoked large EPSCs (middle) or small EPSCs (right). Phostimulations evoked AMPA responses, recorded at −75 mV and blocked with NBQX, and NMDA responses, recorded at +40 mV. The light traces correspond to the sweeps and the average trace is shown in dark. **(F)** EPSCs amplitude evoked by the photostimulation of VMT and EC inputs at −75 mV. Red filled dots correspond to cells responding to VMT photostimulation. Dark blue filled dots correspond to cells displaying large EPSCs whereas light blue filled dots correspond to cells displaying small EPSC to EC photostimulation. Each boxplot shows the 25th, 50th and 75th percentiles and whiskers represent the 5th to the 95th percentile. The median value of EPSC amplitude is smaller for VMT photo-activation (n=10 cells) than for EC photo-activation (n=11 cells; Mann-Whitney test, p=0.0610). **(G)** Epifluorescence microscope images of patched CA1 *slm* GABAergic neuron (green) displaying large EPSCs (left) and small EPSCs (right) to the light stimulation of EC afferents (in red). Scale bars: 250 µm. **(H)** Radial location of patched CA1 *slm* GABAergic neuron displaying large EPSCs and small EPSCs. On the y-axis, 0 corresponds to the *alveus* surface. Dark blue filled dots correspond to cells displaying large EPSCs whereas light blue filled dots correspond to cells displaying small EPSC to EC photostimulation. For each group, the 25th, 50th and 75th percentiles were represented with the same color code. The blue curve represents the average distribution of EC axonal density on the radial axis of CA1 measured on slices corresponding to the small and large EPSC (n=11 slices). Significant radial location difference between large EPSC (n=5 cells) and small EPSC (n=6 cells) was observed (Mann-Whitney, U=2, p=0.0173). **(I)** Same as **(H)** but for disto-proximal location of pached CA1 *slm* GABAergic neuron displaying large EPSCs and small EPSCs. On the y-axis, 0 corresponds to the distal border of CA1 *slm*.The blue curve represents the average distribution of EC axonal density on the disto-proximal axis of CA1 *slm* measured on slices corresponding to the small and large EPSC (n=11 slices). Significant disto-proximal location difference between large EPSC (n=5 cells) and small EPSC (n=6 cells) was observed (Mann-Whitney, U=0, p=0.0043).

In order to describe the connectivity of EC and VMT afferents onto CA1 pyramidal neurons, we next performed patch-clamp recordings from these neurons in the *stratum pyramidale*. As above, recordings were performed in ventral hippocampal slices of P5-7 mice injected with Chronos/tdTomato expression in the EC or VMT (**Figure 4A**). Putative pyramidal neurons were randomly sampled along the radial axis. All cells were filled with biocytin to confirm their identity *post hoc* and recover their axonal and dendritic distribution. The distal dendrites of all morphologically-recovered cells extended into the CA1 *slm* (**Figure 4B**). A much higher fraction of CA1 pyramidal cells was recruited by EC afferents photostimulation as compared to VMT (VMT: 7 %, n=27 cells, EC: 57 %, n=14 cells, Fisher’s exact test, p=0.0012; **Figure 4D1**). The percentage of monosynaptic response across trials was significantly higher when stimulating EC afferents than VMT (VMT: median=0,00 % [iqr=100], n=27 cells, EC: median=55% [iqr=100], n=14 cells, Mann-Whitney test, p=0.0010; **Figure 4D2**). Among the responsive cells, VMT and EC photostimulation evoked EPSCs with similar short delays (VMT: 4.05 ms, n=2 cells, EC: 5.5 ms, n=8 cells; **Figures 4C, S6A**). Overall, VMT afferents photostimulation evoked mainly polysynaptic events in the form of significant increases in EPSCs frequency within one second from the start of the photostimulation (n=12 cells, peak value=0.77, above chance level; **Figures 5A1, 5A2**). The frequency of EPSCs was significantly increased during VMT stimulation (Stim) as compared with the period without stimulation (Stim: mean=8.2 ± 5.4 Hz, No Stim: mean=1.25 ± 0.77 Hz, n=20 cells, Paired t-test, p<0.0001; **Figure 5A3**). Similar increases in frequency were observed when recording Inhibitory Postsynaptic Currents (IPSCs) at the reversal potential for glutamatergic inputs (n=20 cells, peak value=0.74, above chance level, **Figures 5B1, 5B2**; Frequency: Stim: median=12.20 Hz [iqr=26.70], No Stim: median=1.57 Hz [iqr=5.44], n=12 cells, Wilcoxon test, p=0.0005, **Figure 5B3**). These results suggest that the modulation of CA1 pyramidal cells by VMT inputs is mainly polysynaptic and involves local recurrent circuits. Blocking GABAergic transmission significantly decreased EPSCs frequency during the photostimulation period (Ctrl: median=14.80 Hz [iqr=13.50], SR95531: median=3.556 Hz [iqr=5.40], n=7 cells, Wilcoxon test, p=0.0156; **Figures 5C1, 5C2**). Results were confirmed by normalizing events frequency during the stimulation by non-stimulation period frequency (Ctrl: median=10.89 Hz [iqr=11.80], SR95531: median=8.277 Hz [iqr=9.35], n=7 cells, Wilcoxon test, p=0.0469; **Figures S6B, S6C**). Altogether we conclude that VMT inputs are mainly activating GABAergic *slm* neurons which in turn produce, in slices, polysynaptic excitatory events in CA1 pyramids. This is in stark contrast with the monosynaptic EC drive onto CA1 pyramidal neurons.

**Figure 4.**
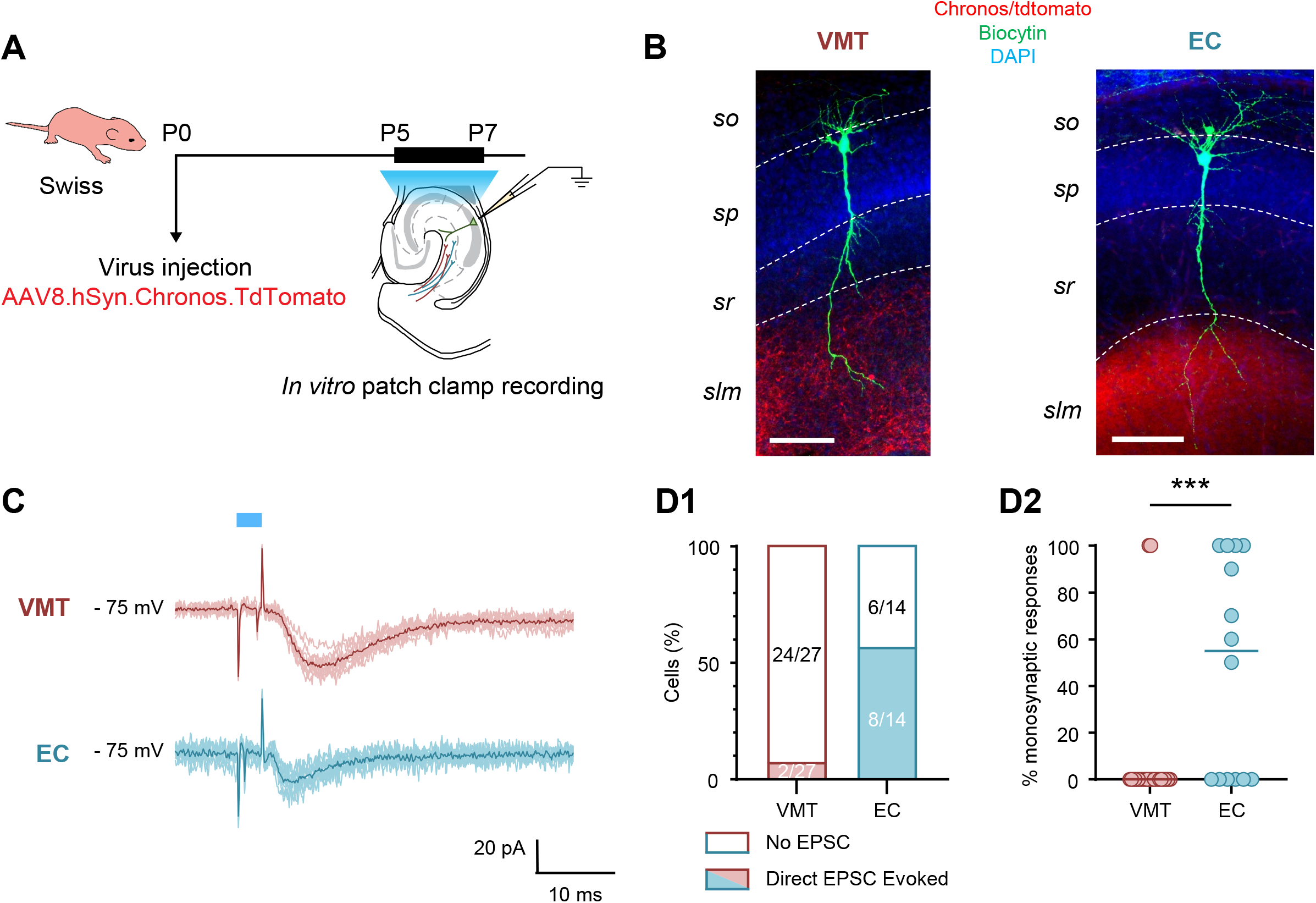
The entorhinal cortex directly recruits CA1 pyramidal neurons. **(A)** Schematic representation of the experimental timeline. **(B)** Dendrites of patched pyramidal neuron (green) reach the CA1 *slm*, where the VMT (left panel) and EC (right panel) axons expressing the Chronos/tdTomato project. Scale bars: 100 µm. **(C)** Monosynaptic glutamatergic responses evoked by VMT (top, red) and EC (bottom, blue) LED photostimulation (3 ms light pulse, blue line). The light traces correspond to the sweeps and the average trace is shown in dark. **(D)** D1, Proportion of CA1 pyramidal neurons displaying EPSCs upon stimulation of VMT (red, 7,41 %) and EC (blue, 57,14 %) afferents. Empty parts represent cells that did not respond to the stimulation (VMT: 92.59 %, EC : 42.86 %). Significant difference between VTM (n=27 cells) and EC (n=14 cells) was observed (Fisher’s exact test, p=0.0010). D2, Percentage of monosynaptic responses evoked by VMT (red) and EC (blue) photostimulation on CA1 pyramidal cells. Each dot corresponds to a cell, lines represent the median. Significant difference between VTM (n=27 cells) and EC (n=14 cells) was observed (Mann-Whitney test, U=99, p=0.0010).

**Figure 5.**
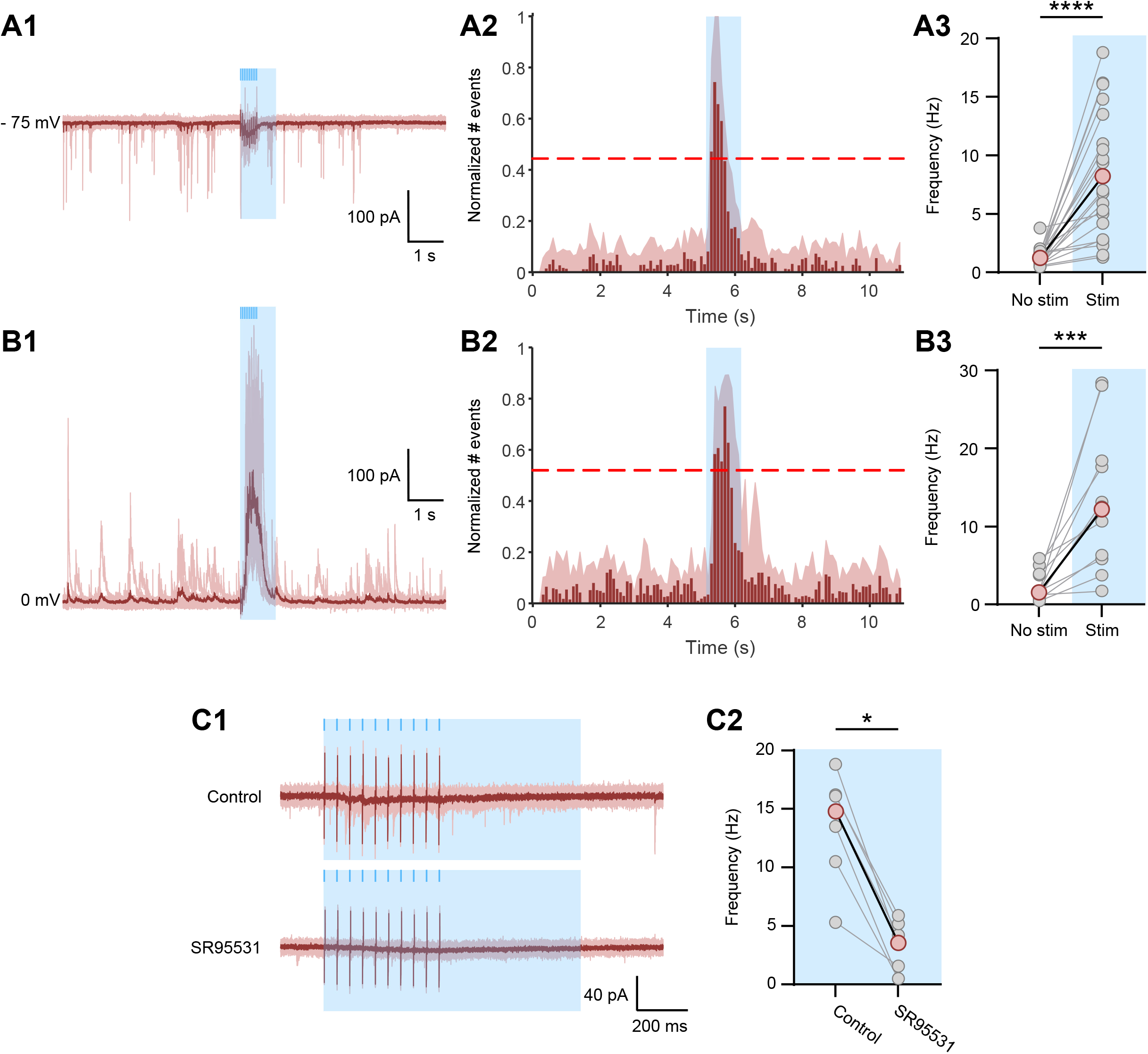
VMT inputs modulate CA1 pyramidal cells through a GABAergic network. **(A)** A1, Spontaneous EPSCs recorded at −75 mV from a CA1 pyramidal cell interrupted by the application of a 10-light pulses train at 20 Hz (3 ms each light pulse, blue lines). The light red traces correspond to the sweeps and the average trace is shown in dark red. The light blue rectangle corresponds to the second following the onset of the first light pulse, named stimulation period (“Stim”). The rest is the non-stimulation period (“No stim”). A2, The normalized number of EPSCs as a function of time represents, for each bin, the median value calculated from the recorded cells. For each cell, the number of EPSCs was summed within 100 ms bins and then normalized by the maximum number of EPSCs observed in a bin. The light red area represents the 75th percentiles for each 100 ms bin. The threshold, in red dotted line, represents the median of the thresholds for each bin, which corresponds to the median value of the 95th percentile calculated from 1000 surrogates. The light blue rectangle corresponds to the “Stim” period defined in 1. A3, EPSCs frequency measured during the No stim and Stim (light blue) periods. Each light gray dot represents a cell, the red dots correspond to the means. Paired comparisons between No stim and Stim periods were tested using the Paired t-test (n=20 cells, t=5.795, p<0.0001). **(B)** Same as **(A)** but for IPSCs. B1, Recording performed at the glutamate reversal potential (0 mV). B2, The normalized number of IPSCs as a function of time was obtained as described in A2. B3, IPSCs frequency measured during the No stim and Stim (light blue) periods. Paired comparisons between No stim and Stim periods were tested using the Wilcoxon test (n=12 cells, W=78.0, p=0.0005). **(C)** 1. EPSCs recorded during the Stim period (light blue rectangle) without (Control) and with GABA_A_R blocker (SR95531). Stim period corresponds to that described in A1. The light red traces correspond to the sweeps and the average trace is shown in dark red. 2. EPSCs frequency measured without (Ctrl) and with GABA_A_R blocker (SR95531) during the Stim periods (light blue). Each light gray dot represents a cell, the red dots correspond to the means. Paired comparisons between Control and SR95531 periods were tested using the Wilcoxon test (n=7 cells, W=-28.0, p=0.0156).

### EC and VMT both exert an excitatory drive onto CA1 circuits in vivo but only EC drives synchronization

We last examined the influence of the two extrinsic inputs on the developing CA1 neuronal activity *in vivo*. To this aim, we injected newborn mice with a virus inducing the expression of the inhibitory DREADD receptor, hM4D(Gi) (AAV9-hSyn-DIO-hM4D(Gi)-mCherry virus), in either the EC or the VMT. For EC expression, we used the Emx1-Cre driver line which specifically targets excitatory cortical neurons. For VMT expression, we used the VGluT2-Cre driver line which specifically targets excitatory subcortical neurons, including thalamic neurons. We imaged the calcium activity in the CA1 pyramidal layer of non-anesthetized pups (P6-8, n=12 pups) using 2-photon excitation of GCaMP6s-expressing neurons (see methods; **Figure 6A**). We only considered experiments for which we confirmed *post hoc* that DREADD-expressing axons reached the *slm* below the imaged area (**Figure S7A**). As a first step, we calibrated the time course of CNO action with extracellular electrophysiological recordings in the CA1 region of Emx1-Cre mouse pups expressing hM4D(Gi) in the hippocampus. A decrease in the number of spikes was observed forty minutes after CNO subcutaneous injection (**Figures S8**). Based on this time course of CNO action, the multineuron calcium activity recorded during the first ten minutes (0-10) of two-photon imaging was compared to the fortieth to fiftieth minute of recording after CNO injection (40-50, **Figure 6A**). The FOVs were the same between 0-10 and 40-50 minutes periods (**Figures 6B, S7C, Supplementary Movies 5-8**), allowing to track the activity of the same neurons. As previously described (Dard et al., 2022; Mohns and Blumberg, 2008; Valeeva et al., 2019), spontaneous neuronal activity observed during the first ten minutes (0-10) alternated between recurring population bursts (Synchronous Calcium Events, SCEs), and periods of low activity (**Figures 6C, 6D, S7D, S7E**). The fraction of cells recruited in SCEs (Dard et al., 2022: median= 0.50 [iqr=0.14], N=13 animals; 0-10: median=0.48 [iqr=0.27], N=12 animals, Mann-Whitney test, p=0.2254; **Figure S7B left panel**) and the transients frequency (Dard et al., 2022: median=1.4 [iqr=1.9], N=13 animals; 0-10: median=1.2 [iqr=2.5], N=12 animals, Mann-Whitney test, p=0.2701; **Figure S7B right panel**) during that initial period were similar to our previous study (Dard et al., 2022). Pharmacogenetic inhibition of VMT or EC resulted in a significant decrease in the frequency of the calcium transients in cells of the pyramidal layer (EC: 0-10: median=0.48 transients/min [iqr=10.91]; 40-50: median=0.10 transients/min [iqr=12.44], n=2522 cells, N=3 animals, Wilcoxon test, W=-1691767, p<0.0001; **Figure 6E top panel**; VMT: 0-10: median=0.77 transients/min [iqr=11.39]; 40-50: median=0.57 transients/min [iqr=15.02], n=1115 cells, N=3 animals, Wilcoxon test, W=-186998, p<0.0001; **Figure 6E bottom panel**). However, the magnitude of EC inhibition was approximately four times greater than that of VMT. We next used a previously developed deep-learning based method (Dard et al., 2022; Denis et al., 2020) to distinguish between putative pyramidal cells (Pyr) and GABAergic neurons of the *so*/*sp* layers (*so*/*sp*INs; **Figures 6F**). EC inhibition decreased the frequency of transients of both cell types (*so*/*sp*INs: median=63% of 0-10 [iqr=430.9], n=164 cells, N=3 animals, One sample Wilcoxon test, p<0.0001; Pyr: median= 32% of 0-10 [iqr=1501], n=1992 cells, N=3 animals, One sample Wilcoxon test, p<0.0001; **Figure 6F top panel**). In contrast, VMT inhibition only affected pyramidal neurons (*so*/*sp*INs: median=99.00% of 0-10 [iqr=5900], n=284 cells, N=3 animals, One sample Wilcoxon test, p=0.7786; Pyr: median= 61.54% of 0-10 [iqr=1400], n=722 cells, N=3 animals, One sample Wilcoxon test, p<0.0001; **Figure 6F bottom panel**). The controls did not show any difference in the frequency of transients (**Figures S7D, S7E, S7F, S7G**). We last analyzed the impact of inhibition of these pathways on SCEs, and found that only EC inhibition would result in a significant decrease in the number of SCEs within 40-50 minutes (EC: n=3 animals; VMT: n=3 animals; Ctrl: n=5 animals; 2-way ANOVA followed by Šidák’s multiple comparisons test, epochs were different for EC inputs, p=0.0021; **Figure 6G**). Since CA1 SCEs are generally preceded by spontaneous motor twitches (Dard et al., 2021; Mohns and Blumberg, 2008; Valeeva et al., 2019), we finally analyzed the pups spontaneous movements but could not find any significant decrease following CNO exposure (**Figure S7H**). Thus, both EC and VMT control pyramidal cell activity in the neonatal CA1, but only EC controls pyramidal cell synchrony driven by motor twitches. This suggests that sensorimotor activity is mainly relayed by the EC to local CA1 circuits.

**Figure 6.**
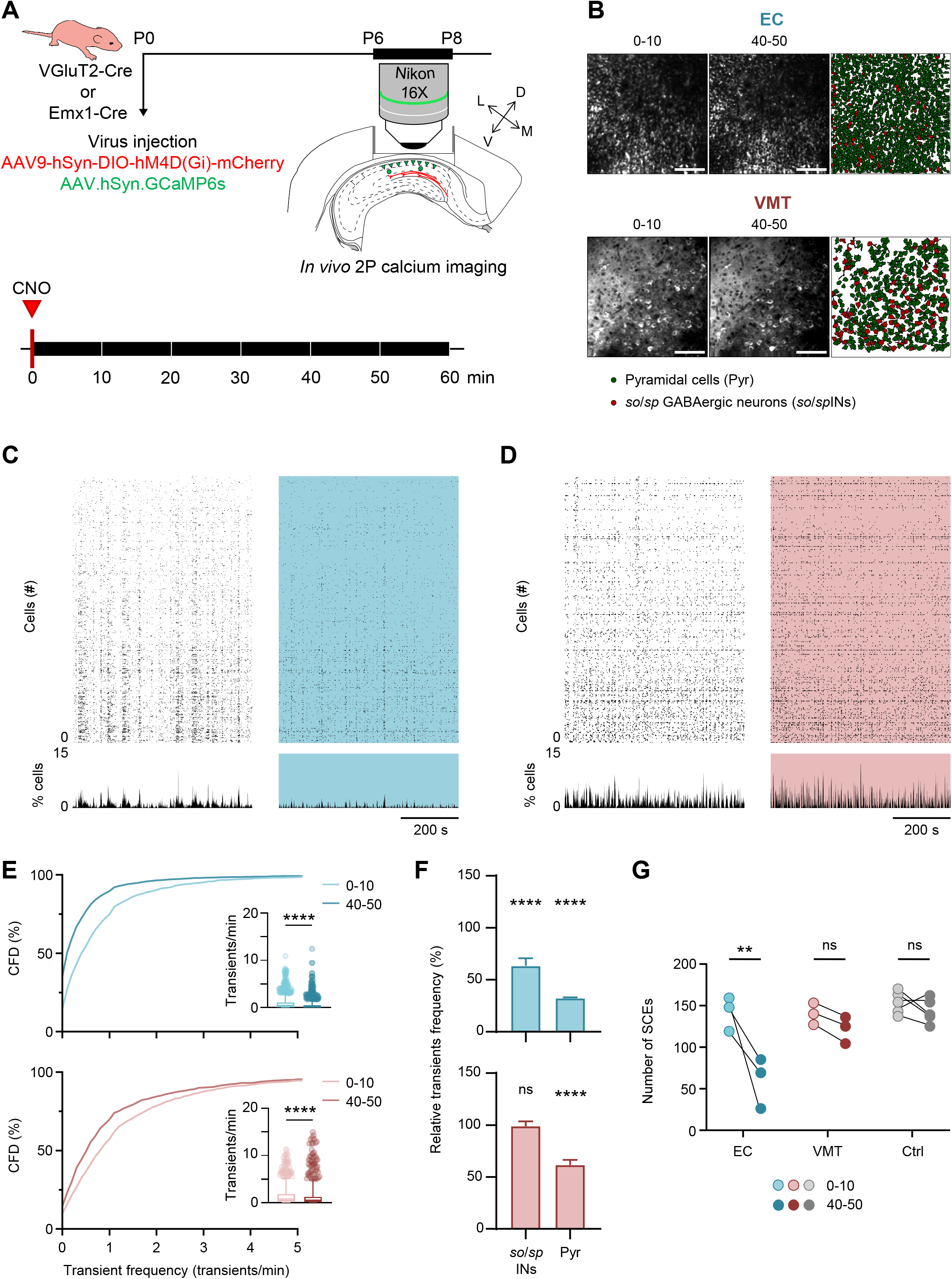
VMT or EC inhibition decreases CA1 network activity in vivo. **(A)** Top: Schematic representation of the experimental timeline. Bottom: Timeline showing the imaging protocol sessions. Several ten-minutes imaging blocks were performed for one hour following the subcutaneous injection of CNO (10 mg/kg). **(B)** FOV of the *stratum pyramidale* imaged during the 0-10 (left) and 40-50 (middle) periods from Emx1-Cre or VGluT2-Cre newborn mouse pups injected, respectively, in the EC or VMT with inhibitory DREADD virus. Scale bars = 100 µm. On the right, the contour map of imaged neurons from the FOV. Inferred *so/sp* GABAergic neurons in red (*so*/*sp*INs), inferred pyramidal cells in green (Pyr). **(C)** Raster plots showing neuronal activity as a function of time during the 0-10 (white) and 40-50 (light blue) periods of Emx1-Cre mouse pups injected in the EC with inhibitory DREADD virus. Graphs below represent the percentage of active cells as a function of time for both periods. **(D)** Same as **(C)** but for DREADD injection in P0 VGluT2-Cre mouse pups’ VMT (light red). **(E)** Cumulative frequency distribution (CFD) plots of the transient frequency during the 0-10 and 40-50 periods for EC (top, n=2522 cells) or VMT (bottom, n=1115 cells) inhibition. The insets show the transient frequency for all cells during 0-10 (light) and 40-50 (dark) epochs. Each boxplot shows the 25th, 50th and 75th percentiles and whiskers represent the 5th to the 95th percentile. The transient frequency was significantly decreased from 40-50 minutes after CNO injection compared to the 0-10 epoch for both EC (Wilcoxon test, W=-1691767, p<0.0001; Kolmogorov-Smirnov test, D=0.2387, p<0.0001) and VMT inhibition (Wilcoxon test, W=-186998, p<0.0001; Kolmogorov-Smirnov test, D=0.1632, p<0.0001). **(F)** Transients frequency measured during 40-50 period in *so*/*sp* GABAergic neurons (*so*/*sp*INs) and pyramidal cells (Pyr) normalized by 0-10 epoch transients frequency for EC (top) or VMT (bottom) inhibition. Each plot shows the 50th and 95th percentile. The median was significantly lower than 100 in *so*/*sp*INs (One sample Wilcoxon test, n=164 cells, theoretical median 100, W=-8140, p<0.0001) and Pyr (One sample Wilcoxon test, n=1992 cells, theoretical median 100, W=-1342710, p<0.0001) for EC inhibition, and significantly lower than 100 in Pyr for VMT inhibition (One sample Wilcoxon test, n=722 cells, theoretical median 100, W=-111959, p<0.0001). **(G)** Number of SCEs measured during 0-10 and 40-50 epochs for EC (N=3 animals) or VMT (N=3 animals) inhibition and control conditions (N=5 animals). Significant difference between 0-10 and 40-50 during EC inhibition was observed (2-way ANOVA followed by Šidák’s multiple comparisons test, inputs-epochs interaction, F(2,8)=7.403, p=0.0151; inputs, F(2,8)=14.32, p=0.0023; epochs, F(1,8)=19.84, p=0.0021; epochs were different for EC inputs, p=0.0021).

## DISCUSSION

Our results demonstrate that early spontaneous activity in CA1 is not only embedded within local CA1 circuits as in the adult (Zutshi et al., 2021), but rather under the combined influence of two extrinsic inputs originating in the VMT and EC. These two inputs are integrated locally through distinct pathways with VMT inputs preferentially targeting *slm* GABAergic neurons.

### Early CA1 dynamics are modulated by extra-hippocampal inputs

Early spontaneous activity in any developing brain region is expected to emerge from the integration of spontaneously active extrinsic inputs originating from more developmentally advanced areas (Donato et al., 2017), onto local circuits capable of self-generating coordinated activity patterns, even when isolated *ex vivo*. Accordingly, we found that both the thalamus and the entorhinal cortex, both older than CA1 -at least for the lateral part of the EC-(Bayer, 1980; Nakagawa, 2019), can modulate the baseline activity and synchronization of local CA1 circuits when inhibited or activated. Activation of these extrinsic inputs is more likely to result in increased network excitability while their inhibition decreases it. Of note, we could not distinguish between LEC and MEC given that targeting the EC at birth with spatial precision already represented an experimental feat. Like the EC inputs (Valeeva et al., 2019), the VMT inputs are spontaneously active in the CA1 region during the first postnatal week. Due to the difficulty to image at a depth of 700 µm combined with the success rate of stereotaxic injections in the VMT at birth, the number of VMT axonal imaging experiments is low and analysis was restricted to quiet rest periods to avoid movement artifacts; in particular we cannot exclude that these inputs could also be activated during spontaneous motor twitches.

The link between the VMT or the EC activity and CA1 dynamics had been previously suggested using simultaneous extracellular electrophysiological recordings in CA1 and the EC or VMT (Hartung et al., 2016; Valeeva et al., 2019). Here, by using chemogenetic tools, we demonstrate a causal link between the activities originating from both inputs and CA1 activity. The time course and strength of the pharmacogenetic inhibition of VMT and EC activity *in vivo* using electrophysiological recordings and performed control experiments where either only CNO or the hM4D(Gi) receptor are provided. Such controls are essential given the possible nonspecific effects of this approach (Roth, 2016). The frequency of single cell calcium transients and of SCEs were used as quantitative readouts of activity. Early sharp waves (eSPWs) and GDPs are likely the electrophysiological correlates of SCEs, measured with calcium imaging *in vivo* and *ex vivo*, respectively, as they are significant network events occurring at similar rates in the same preparations. However, dual simultaneous recordings are needed to confirm this statement.

Our reported effects of VMT or EC inhibition on CA1 dynamics may not only result from direct thalamo-of entorhinal-hippocampal connectivity as both inputs are highly connected to several structures. For example, the VMT innervates the EC as well (Weel and Witter, 1996), and its inhibition could also induce a decrease in the activity of the EC, which would in turn affect the CA1 region. VMT and EC are not the sole extrinsic sources of glutamatergic excitation to CA1. For example, the septum also densely projects to CA1 (Bokor et al., 2002; Kerr et al., 2007), displays an early maturation schedule with a preferential targeting of interneurons (Supèr and Soriano, 1994), conveys proprioceptive inputs and modulates intrinsic CA1 dynamics (Wang et al., 2015; Zhang et al., 2018). However, in contrast to the VMT and the EC, the laminar profile of septal projections does match the excitatory currents driving eSPWs, with their preferential current sink in the *stratum lacunosum moleculare* (*slm)* (Valeeva et al., 2019). The evolution of the current source density profile of eSPWs is interesting. The dual sinks observed in the *stratum radiatum* (*sr*) and *slm* before P5 likely reflect the activation of the entorhino-hippocampal pathway. However, a majority of eSPWs after P5 display a single sink in the *slm* (Valeeva et al., 2019). This may indicate the selective activation of layer 3 MEC neurons from this time point. Whether there are indeed two types of eSPWs with different CSD profiles and associated SCEs depending on their main extra-hippocampal drive remains to be further investigated. Our experiments focused on the period with single sinks in *slm, p*ossibly suggesting a stronger contribution of EC layer III activity in driving SCEs. Future work is needed to study the comparative developmental evolution for the relative weight of VMT and EC inputs to CA1 activity and to describe the in vivo extracellular electrophysiological signature of VMT activity in CA1. Given the protracted development of the mPFC, one possibility could be that the contribution of VMT in driving SCEs *in vivo* is delayed as compared to the EC.

*Ex vivo* experiments allow for the isolation of internally-generated spontaneous activity (GDPs). In these conditions, we found that the photoactivation of the VMT or the EC axons in the CA1 *slm*: (1) can trigger synchronization in the form of GDPs time-locked to the stimulation (in most cases for EC stimulation, in less than half of the cases for VMT stimulation); (2) increased network excitability as revealed by an increase in the rate of GDPs (although in a minority of cases we observed a decrease). The slight difference in the driving of GDPs by VMT and EC likely reflects their integration through different local circuits (see below), as well as their different strength as revealed by the amplitude of the postsynaptic EPSCs in response to stimulation (see also below).

The modulation of CA1 network activity by subcortical inputs from the VMT is reminiscent of the modulation of neocortical bursts by thalamic inputs (Martini et al., 2021; Minlebaev et al., 2011; Mizuno et al., 2018; Molnár et al., 2020). Such bottom-up thalamic drive is integrated within intracortical circuits capable of self-organizing their activity into network bursts, thus producing co-existing network patterns. Interestingly, these thalamic inputs transiently contact local interneurons in the neocortex, the same way VMT inputs are relayed by CA1 *slm* interneurons (Ibrahim et al., 2021; Marques-Smith et al., 2016; Molnár et al., 2020; Tuncdemir et al., 2016).

### Differential intrahippocampal integration of VMT and EC inputs

Our results demonstrate that the VMT and EC inputs integrate different CA1 circuits: (i) their axonal projection distributions differ along the 3 main hippocampal axes; (ii) they display different probabilities to recruit CA1 pyramids; (iii) their *in vivo* network impact diverges.

A striking anatomical segregation of VMT and EC axonal afferents in the CA1 *slm* can be observed using confocal imaging of their respective projections labeled by targeted virus injection at birth. These targeted injections are technically challenging and the site of virus infection may be slightly larger than the anatomical borders of the EC and VMT (**Figures S1, S2**). Still, the labeled axonal tracts are almost exclusively observed in the *slm* of the ventral to intermediate CA1, as expected from anatomical descriptions in the adult (Weel and Witter, 1996). Adding to this differential distribution along the dorso-ventral CA1, VMT and EC afferents segregate along the proximo-distal axis, with VMT targeting the distal *slm* (closer to the subiculum) and EC the proximal *slm* (closer to CA2). This proximo-distal segregation is well described for EC afferents in the adult with LEC projecting distally, like the VMT and MEC proximally (Masurkar et al., 2017, 2020). This may suggest that our stereotaxic injections into the EC were biased towards the MEC at the expenses of the LEC.

This anatomical segregation of projections is functionally reflected in the distribution of the postsynaptic cells responding to their stimulation. Indeed, neurons activated by EC inputs were located in the proximal CA1 *slm*, while neurons responding to VMT inputs were located in the distal *slm*. The probability to observe an EPSC was confirmed by *post hoc* anatomical analysis to report indirectly the number of presynaptic inputs. Therefore the VMT and EC inputs should activate different circuits. There are two major subtypes of neurons in the CA1 *slm*, the glutamatergic Cajal Retzius (CR) cells and the GABAergic interneurons, most of which of the Neurogliaform (NGF) subtype (Anstötz et al., 2015; Chittajallu et al., 2017). Whether different subtypes of *slm* neurons differentially distribute along the proximo-distal axis remains unknown. However, none of the *slm* neurons recorded here and *post hoc* morphologically analyzed displayed the anatomical characteristics of CR neurons whereas putative NGF cells were identified as postsynaptic targets for both VMT and EC inputs. Whether these include the two subtypes of NGF with differential embryonic origins remains to be determined (Overstreet-Wadiche and McBain, 2015). In any case, this identifies NGF cells as central nodes integrating these two major inputs onto the developing CA1. Interestingly, CA1 NGFs are also a major postsynaptic target of CR cells, an important player in CA1 circuit development (Quattrocolo and Maccaferri, 2014). In addition, NGFs were shown to be central in synchronizing interneuron networks using a combination of synaptic and electrical signaling (Overstreet-Wadiche and McBain, 2015; Zsiros et al., 2007). It is therefore possible that NGF neurons in the developing CA1 function in providing an unspecific and broad feedforward activation of local inhibitory circuits in response to extra-hippocampal inputs.

If the proximo-distal distribution of *slm* GABAergic neurons is unknown, less is true for their *stratum pyramidale* glutamatergic counterparts which are known to segregate along this axis according to their birthdate (Cossart and Khazipov, 2021). It is also known that CA1 pyramids, in the adult, are directly contacted by EC inputs, with cells located in deeper portions of the CA1 radial axis more likely to display direct contacts (Masurkar et al., 2017). Given that deep CA1 pyramids are older than superficial ones, these are likely to be more developed with a dendritic arborisation reaching the *slm* (Tyzio et al., 1999). Morphological analysis of the CA1 pyramids displaying evoked EPSCs following EC input photoactivation may indicate whether these share similar dendritic properties and soma location. If more than half of the recorded CA1 pyramids displayed EC-evoked EPSCs, almost none was directly stimulated by VMT photoactivation. The possibility that VMT evoked EPSCs would be undetected at the soma due to their distal origin is unlikely given that EC inputs are detected (despite a similar distal origin) and because of the inherent electrotonic compactness of immature neurons. Interestingly, the absence of direct VMT inputs onto most pyramids was also reported in the adult CA1 (Andrianova et al., 2021), indicating that this different innervation does not reflect a protracted maturation of VMT axons onto CA1 pyramids (even though VMT CA1 inputs arrive three days later than EC inputs (Supèr and Soriano, 1994)). Whether the rare CA1 pyramids we found to be directly activated by VMT inputs in the developing CA1 reflect a transient developmental connection or reveal a rare subtype of CA1 pyramidal neuron remains to be further studied.

The differential anatomical and functional connectivity of EC and VMT inputs is reflected in the single-cell *in vivo* response to their chemogenetic inhibition. Of note, these single-cell responses were principally monitored in the *stratum pyramidale*, away from the initial site of VMT and EC input integration. As discussed above, EC inhibition results in a decrease in the fraction of active neurons involved in SCEs, an effect that is not observed when inhibiting the VMT, which only reduces the fraction of pyramidal cell calcium transients. The differential impact of VMT versus EC onto CA1 SCEs likely indicates the EC as the main relay station for sensorimotor inputs generated by spontaneous motor twitches (Karlsson et al., 2006; Rio-Bermudez and Blumberg, 2021; Valeeva et al., 2019) and is supported by a larger excitatory drive from EC onto CA1 principal cells. The lack of VMT modulation of interneuron activity in the CA1 *stratum pyramidale in vivo* contrasts with the generation of evoked polysynaptic GABAergic events and GDPs by activation of VMT inputs *ex vivo*. This may have several explanations, including the possible lack of direct synaptic inputs from CA1 slm interneurons and CA1 pyramids onto *sp* interneurons at these early stages (Dard et al., 2022). However, *ex vivo* experiments indicate that *slm* interneurons are rather well connected into local circuits since their optogenetic activation alone induces polysynaptic GABAergic responses in principal cells. Such ability to trigger synchrony through the selective activation of interneuron circuits may be related to the excitatory action of the transmitter in slices (Dzhala et al., 2012; Valeeva et al., 2016). The decrease of CA1 principal cell activity following VMT *in vivo* inhibition could also be explained by a direct excitatory GABAergic input, for example from NGF cells onto pyramids (but see (Murata and Colonnese, 2020)) or by a local disinhibitory circuit which remains to be elucidated.

In conclusion, our study shows how bottom-up inputs (transmitted from the EC) and higher order thalamic inputs (transmitting top-down information from the mPFC in the adult (Ferraris et al., 2021)), are differentially integrated within local intermediate to ventral CA1 circuits and modulate or even drive their early dynamics. If these inputs segregate between CA1 pyramids, they converge onto local *slm* GABAergic interneurons, including NGF cells of the *slm*. This is reminiscent of the developing neocortex, where layer 1 NGF neurons are first driven by bottom-up inputs, which in turn regulate the establishment of top-down connections from integrative areas (Che et al., 2018; Ibrahim et al., 2021). Whether similar mechanisms operate in the developing CA1 opens an interesting venue for research, this even more given the role of the nucleus reuniens on developmental brain disorders implicating impaired cognitive functions (Cassel et al., 2021; Ferraris et al., 2021; Weel and Witter, 2020).

## ACKNOWLEDGMENTS

This work was supported by the European Research Council under the European Union’s, Horizon 2020 research and innovation program Grant #646925 and #951330 (HOPE), the Fondation Bettencourt Schueller. E.L. was funded by the “Ministère de l’Enseignement Supérieur, de la Recherche et de l’Innovation”, A*Midex foundation and the French National Research Agency funded by the French Government « Investissements d’Avenir » program (NeuroSchool, nEURo*AMU, ANR17-EURE-0029 grant), by the Fondation Bettencourt Schueller and is currently supported by ERC. R.F.D. was funded by the “Ministère de l’Enseignement Supérieur, de la Recherche et de l’Innovation”, and currently by the Fondation pour la Recherche Médicale (grant No. FDT202106012824). S.M. is funded by the “Ministère de l’Enseignement Supérieur, de la Recherche et de l’Innovation”. C.F. is funded by ERC. M.B. was funded by the Fyssen Foundation, the Fondation pour la Recherche Médicale (grant No. SPF20170938593), and by the European Union (Marie Skłodowska-Curie individual fellowship, grant No. 794861—IF-2017). We would like to thank V. Crepel and A. Represa for sharing equipment, F. Michel from the INMED imaging facility (InMagic) and INMED animal facility staff. We are grateful to M. Esclapez and A. de Chevigny for providing, respectively, the VGlut2-Cre and the Emx1-Cre mouse lines.

## AUTHORS CONTRIBUTION

EL, AB, and RC designed research. EL performed stereotaxic viral injection and RFD performed intraventricular viral injections. RFD performed surgery for *in vivo* 2-photon calcium imaging experiments and RFD, EL and MAP performed 2-photon calcium imaging acquisition. EL performed *ex vivo* 2-photon calcium imaging and patch-clamp recordings experiments. MB supervised *ex vivo* patch-clamp recording experiments. EL and RFD performed 2-photon calcium imaging analysis and EL performed *ex vivo* patch-clamp recordings analysis. SM and PPLS performed *in vivo* extracellular electrophysiological recordings and their analysis. CF, MG-K, AB, and EL performed histology experiments. EL, AB and CF performed histological analysis. RC and EL wrote the paper.

---

**Figure S1.**
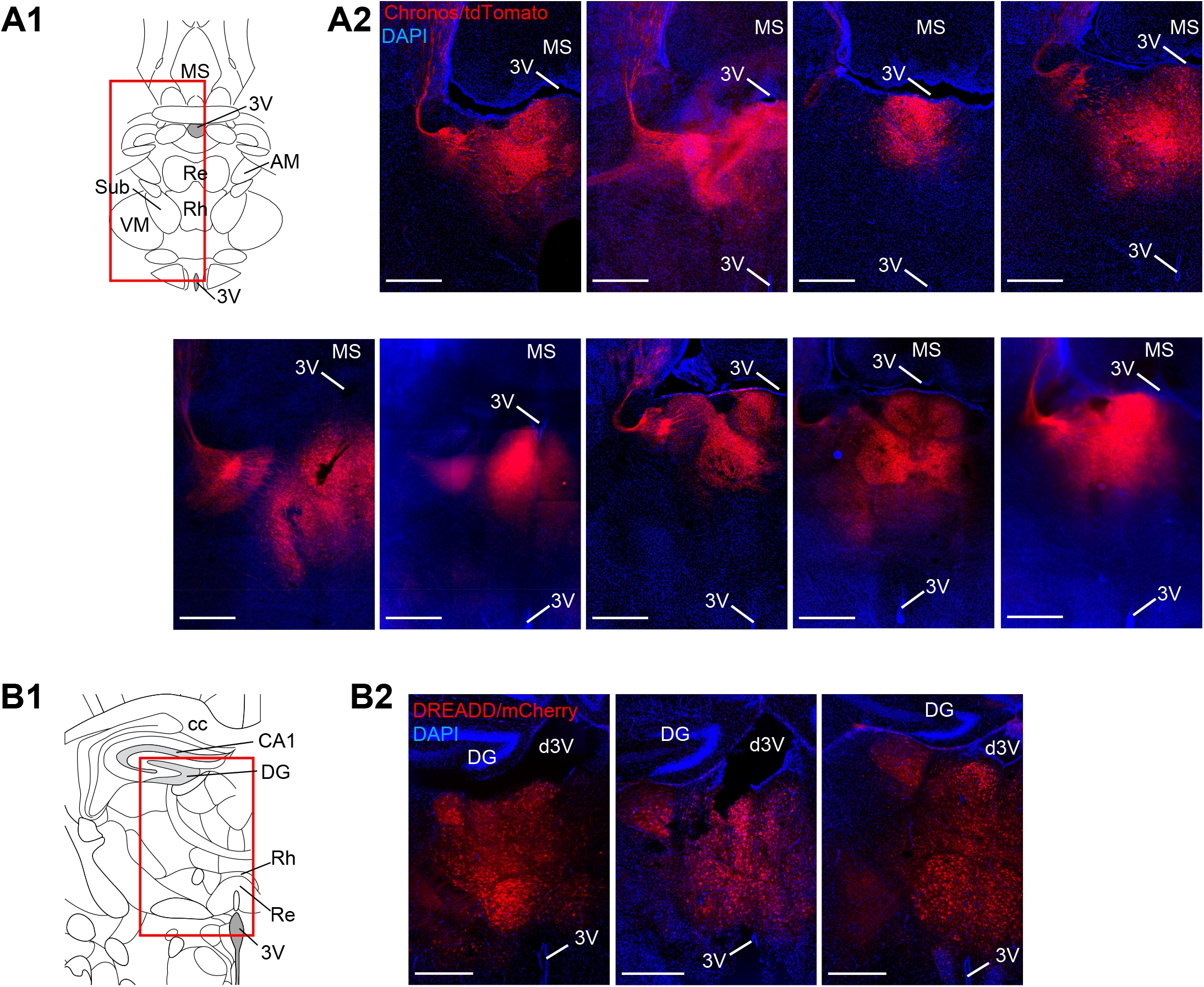
Examples of VMT injection sites. **(A)** A1, Schematic representation of the ventral midline thalamus (VMT) from an horizontal slice adapted from (Franklin and Paxinos, 2007). The red rectangle corresponds to the area of the confocal images in Figure A2. A2, Examples of confocal or epifluorescence microscope images of P5-P7 brain horizontal sections from different wild-type SWISS mouse pups injected at P0 with Chronos/tdTomato virus in the VMT, composed by the reuniens nucleus (Re) and the rhomboid nucleus (Rh). Chronos/tdTomato virus was not expressed in the medial septum (MS). Scale bars: 500 µm **(B)** B1, Schematic representation of the medial part of the thalamus from a coronal slice adapted from (Paxinos et al., 2020). The red rectangle corresponds to the area of the confocal images in Figure B2. B2, Confocal or epifluorescence microscope images of P6-P8 brain coronal sections from VGluT2-Cre mouse pups (N=3 animals) injected at P0 with DREADD/mCherry virus in the VMT. These animals correspond to the ones used during *in vivo* 2-photon calcium imaging experiments. Injection site is located between the 2 parts of the 3rd ventricle (3V). DREADD/mCherry virus was not expressed in the dentate gyrus of the hippocampus (DG). Scale bars: 500 µm

**Figure S2.**
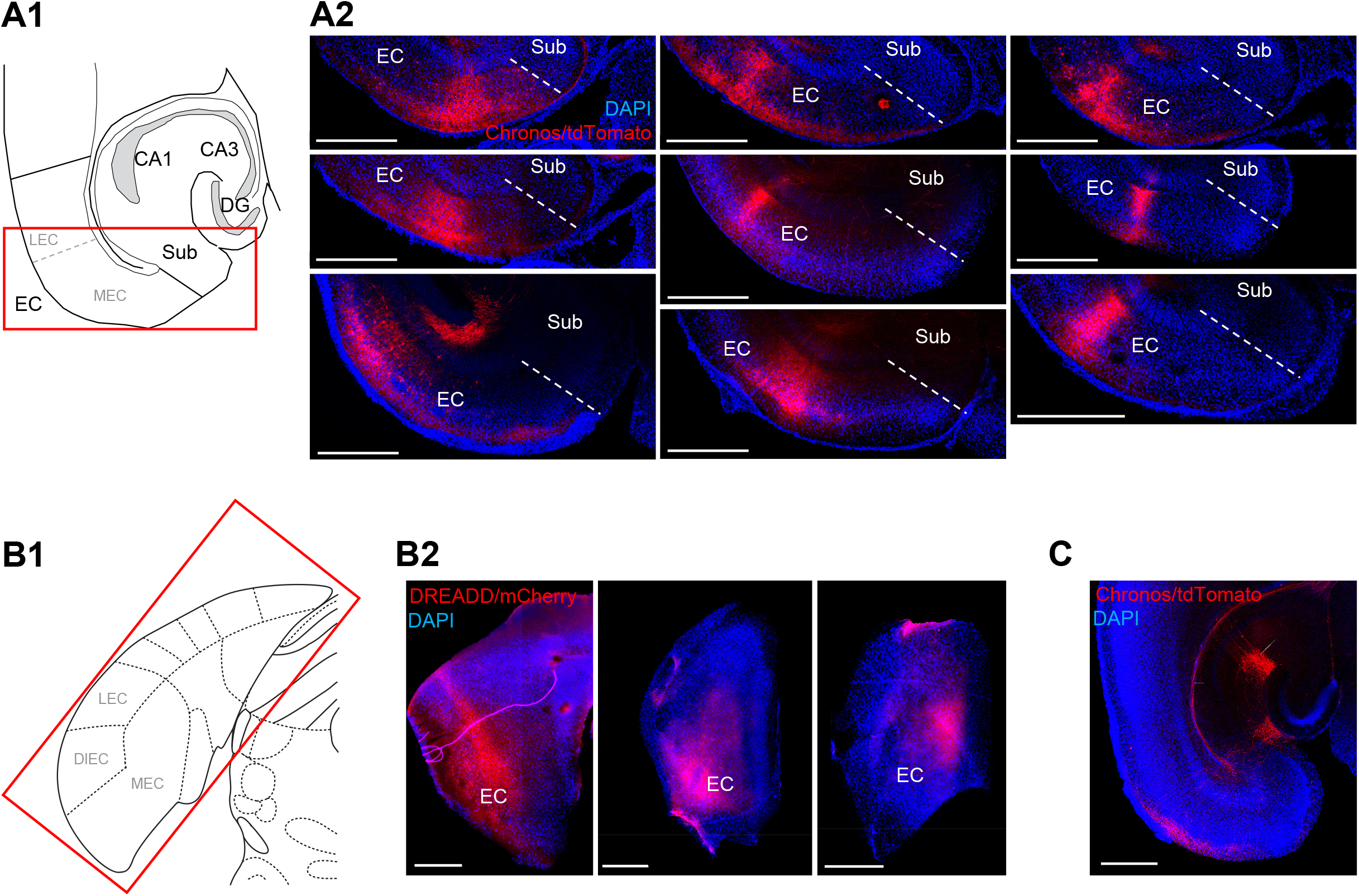
Examples of EC injection sites. **(A)** A1, Schematic representation of the entorhinal cortex (EC) on an horizontal slice adapted from (Franklin and Paxinos, 2007). The red rectangle corresponds to the area of the confocal images in Figure A2. A2, Examples of confocal or epifluorescence microscope images of P5-P7 brain horizontal sections from different wild-type SWISS mouse pups injected at P0 with Chronos/tdTomato virus in the EC, composed by medial entorhinal cortex (MEC) and the lateral entorhinal cortex (LEC). Chronos/tdTomato virus was not expressed in the subiculum (Sub). Scale bars: 500 µm **(B)** B1, Schematic representation of the EC on a coronal slice adapted from (Paxinos et al., 2020). The red rectangle corresponds to the area of the confocal images in Figure B2. B2, Confocal or epifluorescence microscope images of P6-P8 brain coronal sections from Emx1-Cre mouse pups (N=3 animals) injected at P0 with DREADD/mCherry virus in the EC, composed by the MEC, the LEC and the dorsal intermediate entorhinal cortex (DIEC). These animals correspond to the ones used during *in vivo* 2-photon calcium imaging experiments. Scale bars: 500 µm **(C)** EC fibers expressing Chronos/tdTomato virus project to the contralateral hippocampus. EC fibers reached the contralateral hippocampus through the fimbria and then the alveus (arrow). Fibers are crossing radially CA1 region (arrowhead) to reach CA1 *slm* (double arrowhead).

**Figure S3.**
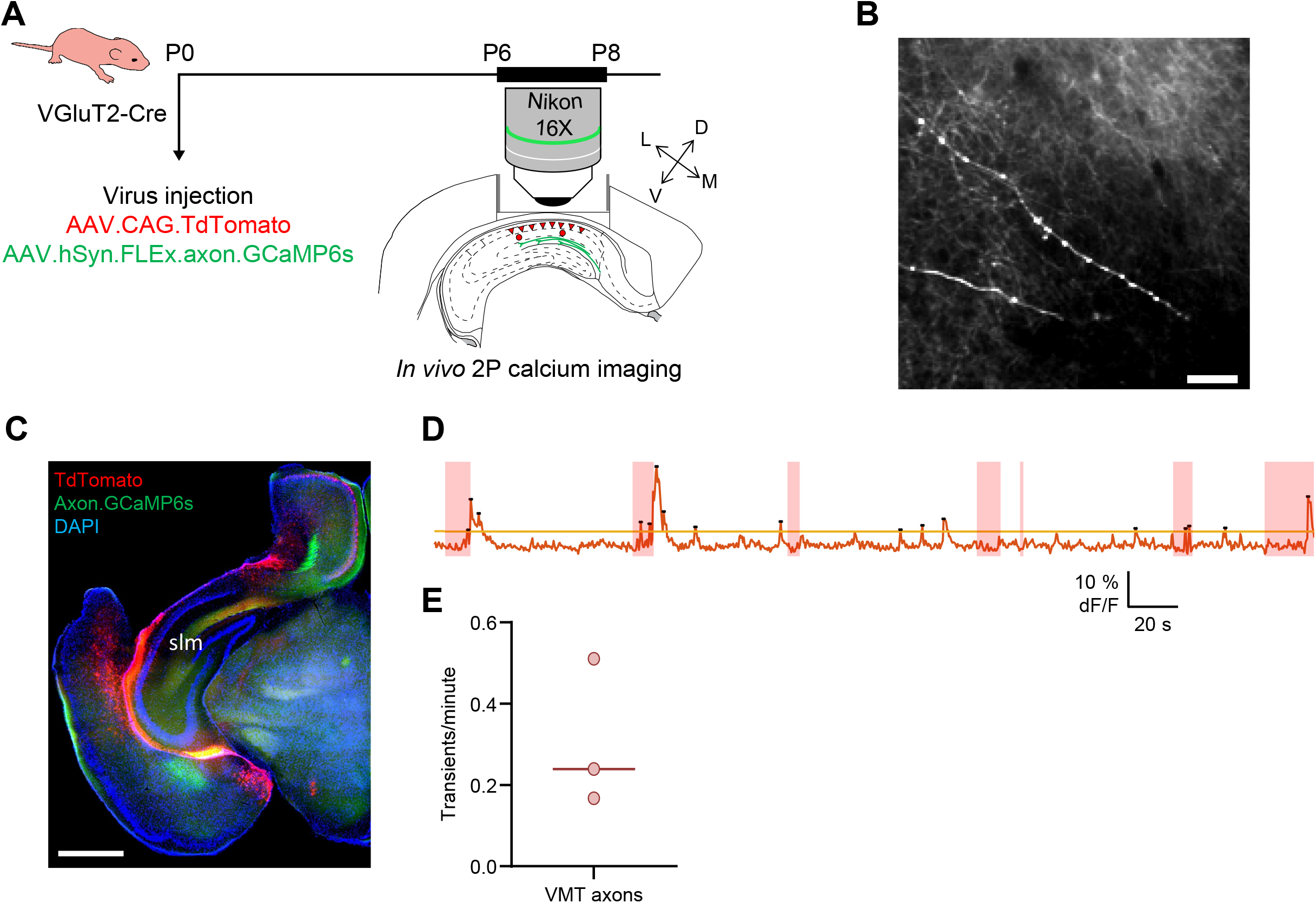
In vivo 2-photon calcium imaging of VMT axons in the CA1 slm at the end of the first postnatal week. **(A)** Schematic representation of the experimental timeline. **(B)** Averaged image from the imaged FOV in the CA1 *slm* of a VGluT2-Cre mouse injected with Cre-dependent axon-GCaMP6s virus at P0 in the VMT. Scale bars = 50 µm. **(C)** Coronal sections showing the location of the cannula and the glass window above the intermediate hippocampus. The VMT fibers (green) were present within the field of view (FOV) in the *slm* of CA1. Scale bars: 1 mm. **(D)** Example of a denoised dF/F signal extracted from an active VMT axonal branch. Light red rectangles correspond to the z-movement periods. The threshold corresponds to the median of the free z-movements denoised trace plus 3 times its interquartile range. Black dots correspond to the significant peaks. Significant transients were conserved if their onset was out of a movement period. **(E)** Distribution of the number of transients per minute measured from active VMT branches expressing GCaMP6s in P6-P8 mouse pups. Each dot corresponds to a FOV (n=3 FOV, N=2 animals), lines represent the median.

**Figure S4.**
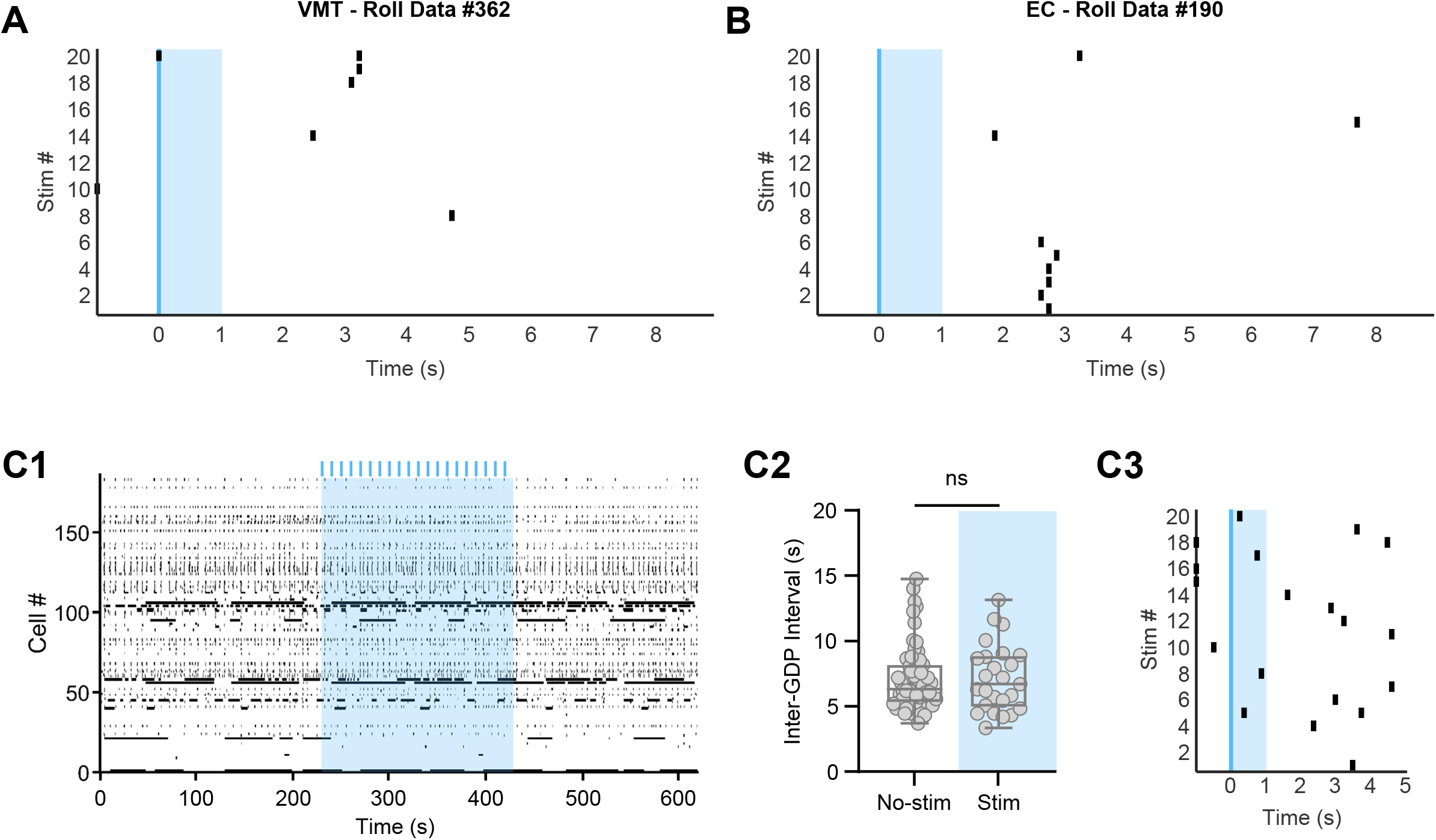
Control analysis and experiment for *ex vivo* 2-photon calcium imaging experiments. **(A)** Example of a surrogate obtained by rolling the time stamp of the GDPs detected during the analysis of an *ex vivo* 2-photon calcium imaging movie to determine the threshold beyond which the association of GDPs with VMT stimulation exceeds the chance level in this movie. **(B)** Same as **(A)** but for EC stimulation. **(C)** Example of control (Ctrl) photostimulation effect on the network activity of developing CA1 observed in slice. The Ctrl experiment consisted of performing stimulation protocols on slices that didn’t express opsin. C1, Raster plots showing neuronal activity as a function of time. The light blue rectangle corresponds to the photostimulation period during which phasic stimulations at 0.1 or 0.2 Hz were applied (blue lines). C2, Measurement of the inter-GDP interval during the period of photostimulation (light blue) and non-stimulation corresponding to the raster plot shown in 1. Each dot corresponds to the measured IGIs during non-stimulation (“No-Stim”) and stimulation (“Stim”) periods. Each boxplot shows the 25th, 50th and 75th percentiles and whiskers represent the 5th to the 95th percentile. No significant difference between. No-Stim and Stim periods was observed (No-Stim: median=6.324 s [iqr=11.04], n=59 IGIs, Stim : median=6.696 s [iqr=9.796], n=25 IGIs, Mann-Whitney, U=725.5, p=0.9091). C3, GDPs distribution during the stimulation period that correspond to the raster plot shown in 1. GDP occurrence (black rectangles) as a function of time following 0.1 Hz phasic Ctrl stimulations. Twenty consecutive light stimulations were centered to time 0 (blue line). Among them, 4 stimuli were followed by GDP in the following second (light blue rectangle, 99^th^ percentile=6).

**Supplementary Table 1 (related to Figure 4).**
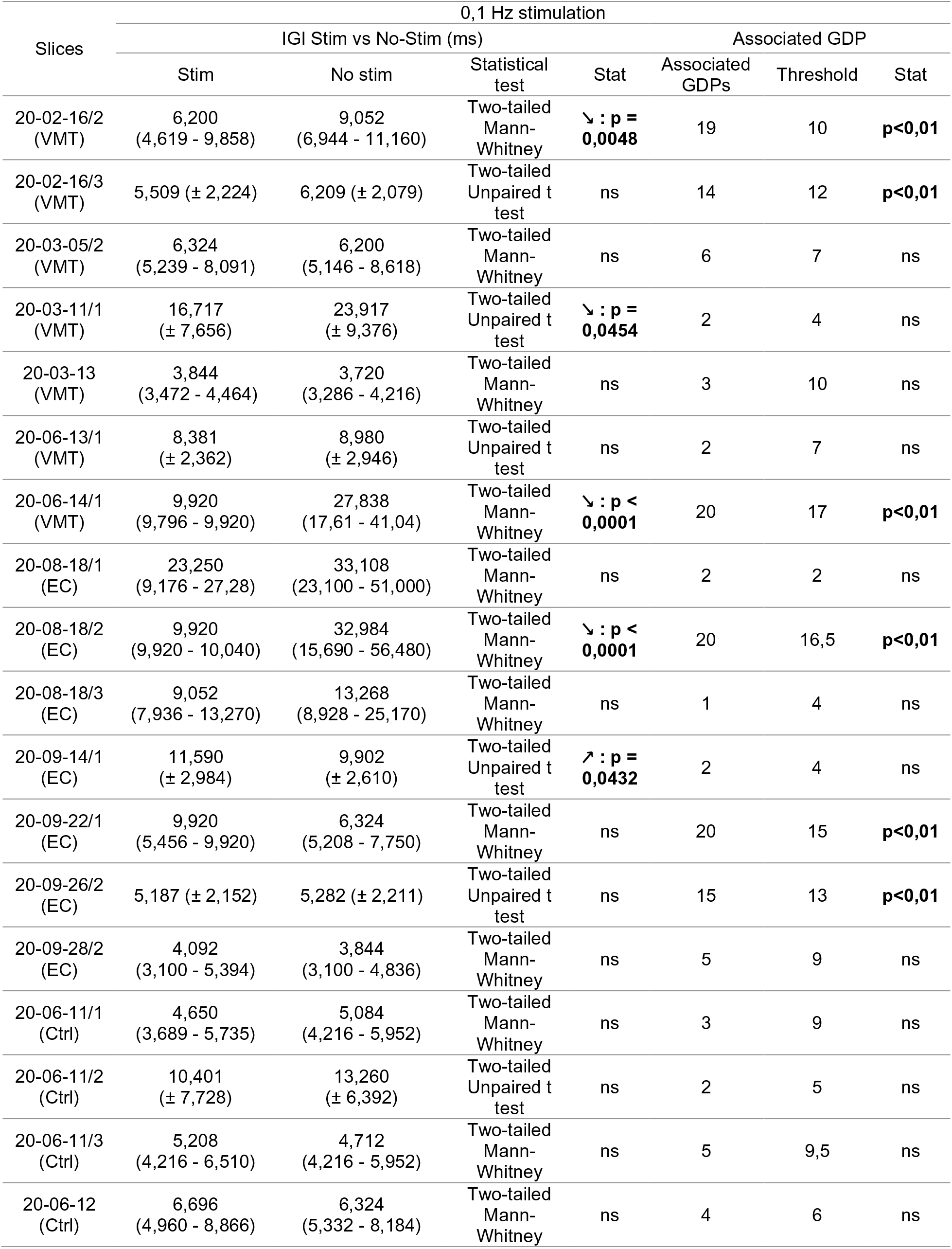
Effect of VMT, EC and Ctrl photostimulation at 0,1 Hz on the IGI and the GDP association to the stimulation. Upward pointing arrow means significant increase of IGI; downward pointing arrows mean decrease of IGI. The IGI values correspond to the median (25th percentile - 75th percentile) or the mean (± SD).

**Supplementary Table 2 (related to Figure 4).**
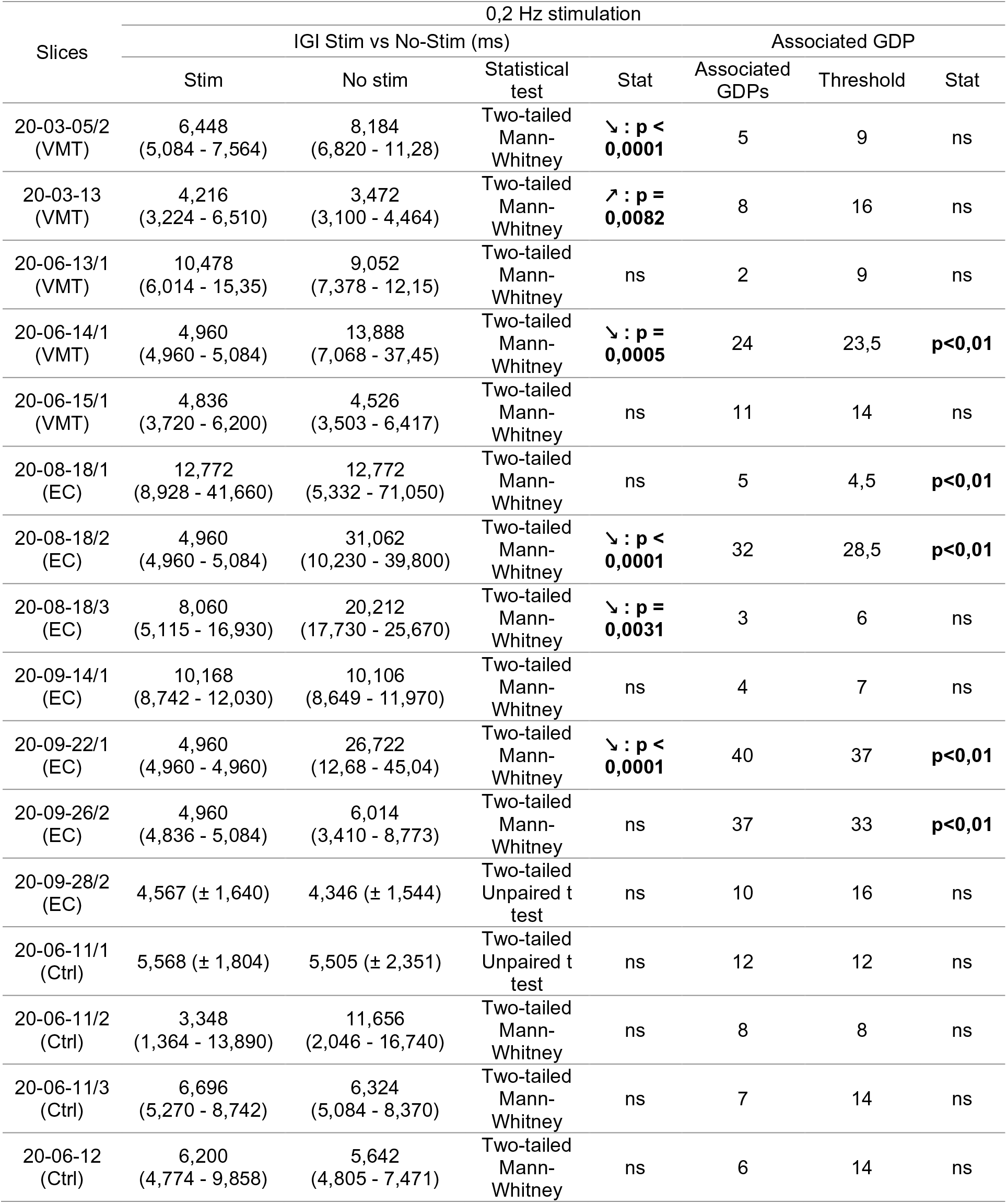
Effect of VMT, EC and Ctrl photostimulation at 0,2 Hz on the IGI and the GDP association to the stimulation. Upward pointing arrow means significant increase of IGI; downward pointing arrows mean decrease of IGI. The IGI values correspond to the median (25th percentile - 75th percentile) or the mean (± SD).

**Figure S5.**
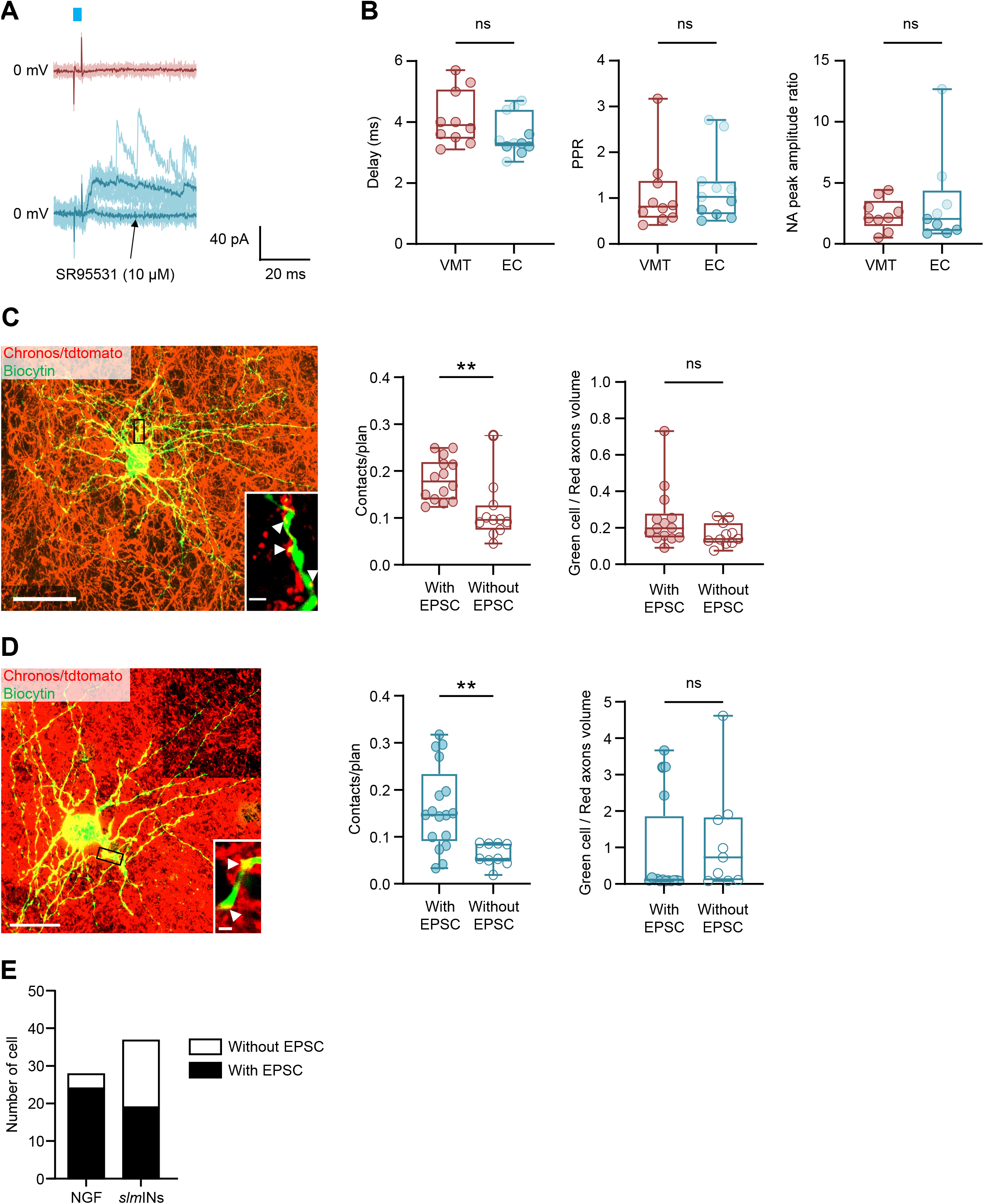
Supplementary electrophysiological and anatomical properties of patched CA1 *slm* GABAergic neurons. **(A)** Monosynaptic GABAergic responses evoked on CA1 *slm* GABAergic neurons, at the EPSCs reversal potential (0 mV), by EC (bottom, blue) but not by VMT (top, red) LED photostimulation (3 ms light pulse, blue rectangle). The monosynaptic GABAergic responses evoked by EC photostimulation were blocked by GABA_A_R inhibitor, SR95531. The light traces correspond to the sweeps and the average trace is shown in dark. **(B)** From the left to the right: delay, paired pulse ratio (PPR), and NMDA/AMPA (N/A) peak amplitude ratio measured on CA1 *slm* GABAergic neurons after photostimulation of VMT or EC inputs. PPR was measured by two 3-ms light pulses at 20 Hz. Red filled dots correspond to cells responding to VMT photostimulation. Dark blue filled dots correspond to cells displaying large EPSCs whereas light blue filled dots correspond to cells displaying small EPSC to EC photostimulation. Each boxplot shows the 25th, 50th and 75th percentiles and whiskers represent the 5th to the 95th percentile. No significant difference between VMT (n=10 cells) and EC (n=11 cells) was observed for the delay (Mann-Whitney test, U=31,50, p=0.1017) and the PPR (Mann-Whitney test, U=46, p=0.5573). No significant difference between VMT (n=9 cells) and EC (n=9 cells) was observed either for N/A peak amplitude ratio (Mann-Whitney test, U=39, p=0.9314). **(C)** Left panel: representative maximum intensity projections of confocal z-stacks showing innervation of a patched CA1 *slm* GABAergic neuron by VMT afferents. Inset: single optical sections (thickness 0.4 µm) showing magnifications of contacts (arrow heads) between a patched neuron dendrite (green) and fibers from the VMT (red). Scale bars: 20 µm for the principal image; 2 µm for the insert. Middle panel: Putative contacts number per plan between patch CA1 *slm* GABAergic neurons and VMT axons. Filled dots correspond to responding cells and empty dots correspond to no-responding cells. Each boxplot shows the 25th, 50th and 75th percentiles and whiskers represent the 5th to the 95th percentile. Significant difference between responding cells (“With EPSC”, n=14 cells) and no-responding cells (“Without EPSC”, n=11 cells) was observed (Mann-Whitney, U=21, p=0.0014). Right panel: Patched cell volume / VMT axons volume ratio compared between responding cells (With EPSCs) and no responding cells (Without EPSCs). Filled dots correspond to responding cells and empty dots correspond to no-responding cells. Each boxplot shows the 25th, 50th and 75th percentiles and whiskers represent the 5th to the 95th percentile. No significant difference between responding cells (With EPSC, n=14 cells) and no-responding cells (Without EPSC, n=11 cells) was observed (Mann-Whitney, U=46, p=0.0954). **(D)** Same as **(C)** but for Chronos/tdTomato injection in P0 SWISS mouse pups’ entorhinal cortex. Scale bars: 20 µm for the principal image; 2 µm for the insert. Significant difference between responding cells (“With EPSC”, n=17 cells) and no-responding cells (“Without EPSC”, n=9 cells) was observed for the number of contacts (Mann-Whitney, U=24, p=0.0035). No significant difference between responding cells (With EPSC, n=16 cells) and no-responding cells (Without EPSC, n=9 cells) was observed for the patched cell volume / EC axons volume ratio (Mann-Whitney, U=53, p=0.3014). **(E)** Proportion of putative neurogliaform-like (NGF) and non-neurogliaform-like (*slm*INs) cells that received or not monosynaptic contacts from the VMT and the EC fibers present in the CA1 *slm*. The proportion of NGF responding to the photostimulation was significantly greater than that of *slm*INs (NGF: n=28 cells, With EPSC=85.71 %, Without EPSC=14.29 %; *slm*INs: n=37 cells, With EPSC=51.35 %, Without EPSC=48.65 %; Fisher’s exact test, p=0.0042).

**Figure S6.**
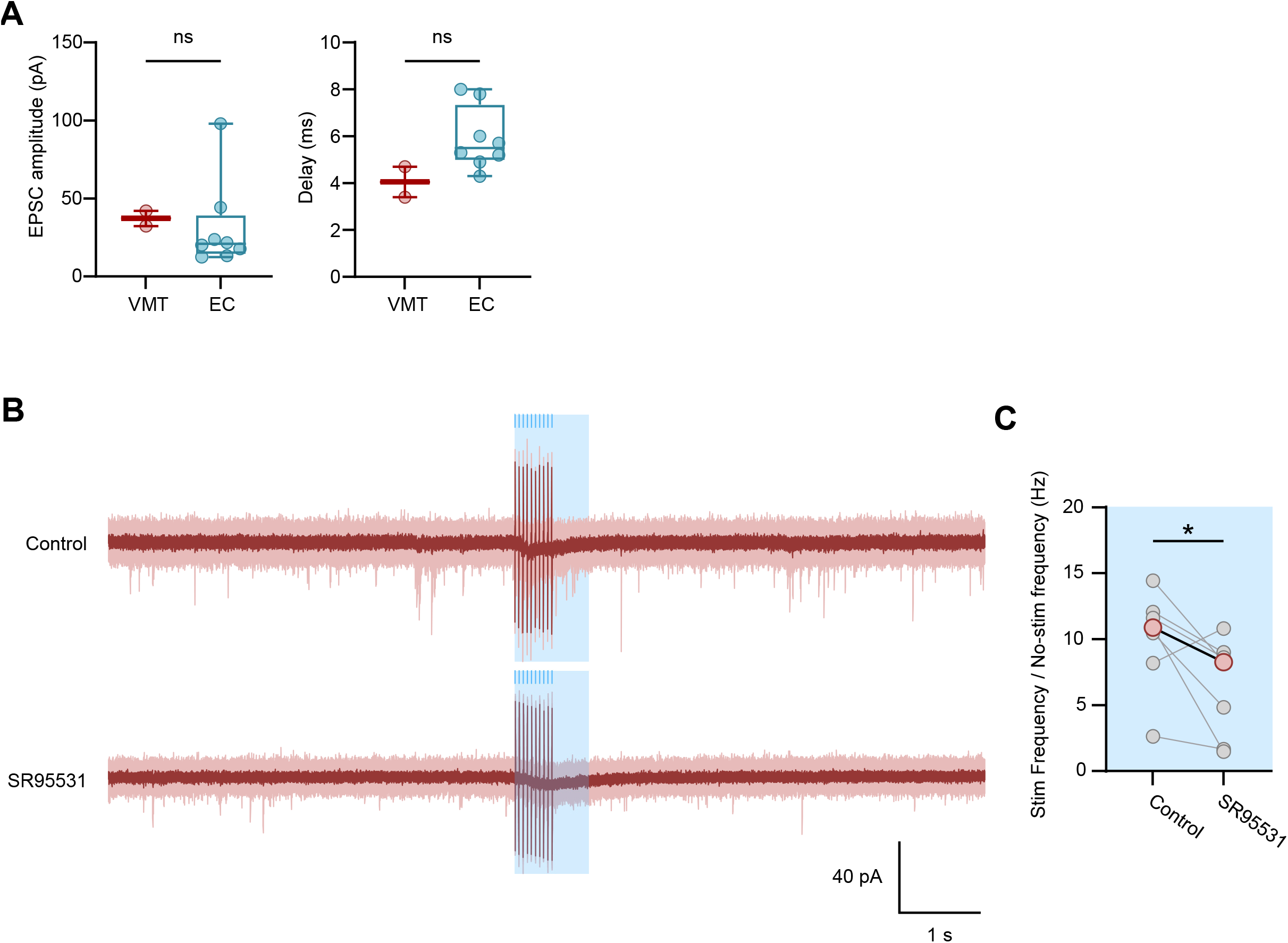
Supplementary electrophysiological properties of patched CA1 pyramidal neurons during stimulation of the VMT inputs. **(A)** From the left to the right: Amplitude and delay measured on CA1 pyramidal neurons that respond to the photostimulation of VMT or EC inputs. Red filled dots correspond to cells responding to VMT photostimulation and blue filled dots correspond to EC photostimulation. Each boxplot shows the 25th, 50th and 75th percentiles and whiskers represent the 5th to the 95th percentile. **(B)** Top panel: Spontaneous EPSCs recorded without drug at −75 mV from a CA1 pyramidal cell interrupted by the application of a 10-light pulses train at 20 Hz (3 ms each light pulse, blue lines). The light red traces correspond to the sweeps and the average trace is shown in dark red. The light blue rectangle corresponds to the second following the onset of the first light pulse, named stimulation period (“Stim”). The rest is the non-stimulation period (“No stim”). Bottom panel: Same as the top panel but after application of the GABA_A_R blocker SR95531. **(C)** Stimulation period EPSCs frequency normalized by the no-stimulation period EPSCs frequency measured without (Ctrl) and with GABA_A_R blocker (SR95531). Each light grey dot represents a cell, the red dots correspond to the means. Paired comparisons between Control and SR95531 periods were tested using the Wilcoxon test (n=7 cells, W=-24.0, p=0.0469).

**Figure S7.**
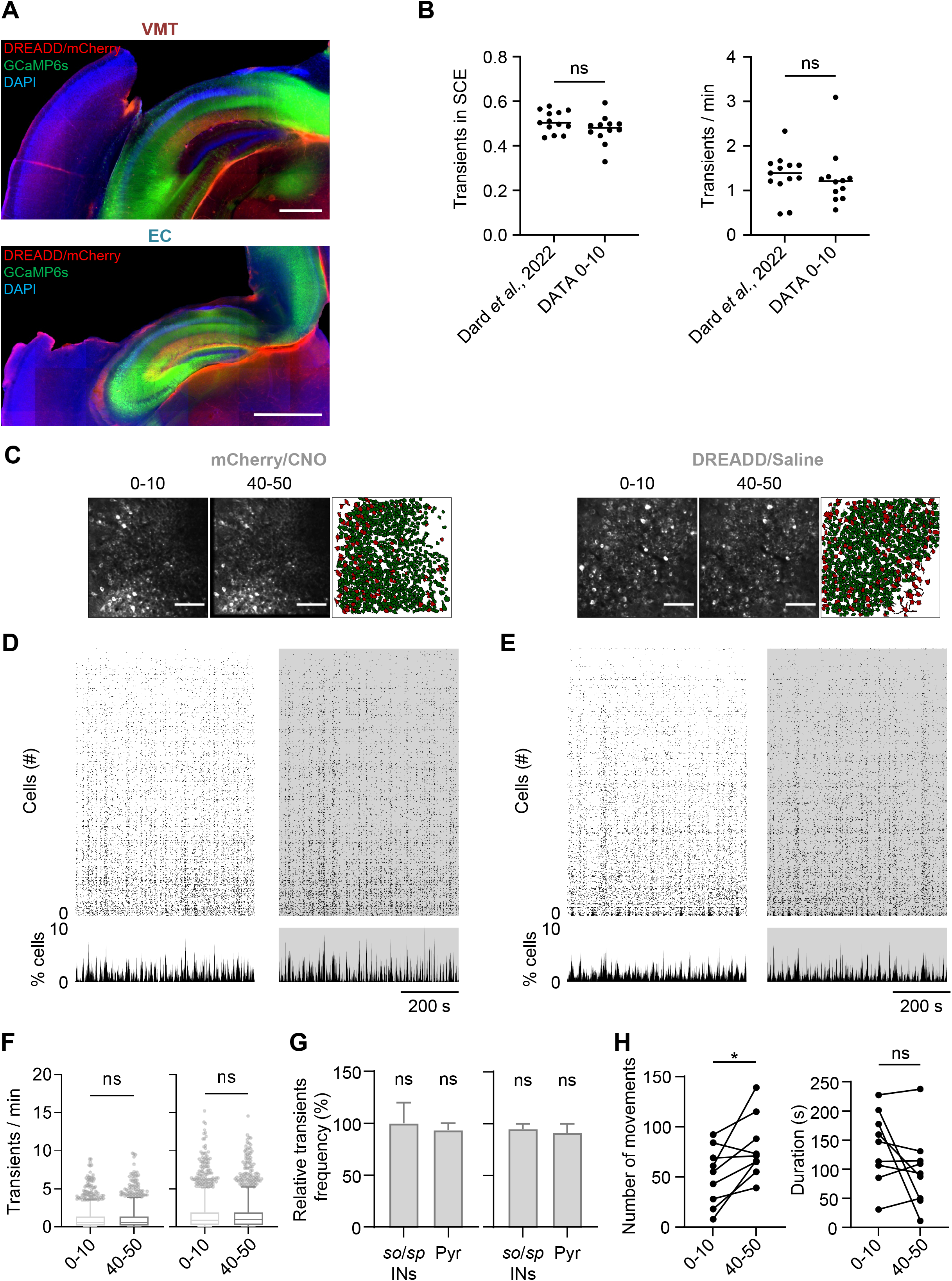
Controls for *in vivo* inhibition experiments with the DREADD receptor. **(A)** Coronal sections showing the location of the cannula and the glass window above the intermediate hippocampus. The VMT (top) and EC (bottom) fibers, in red, were also present under the field of view (FOV) in the *slm* of CA1. Scale bars: 500 µm. **(B)** The ratio of calcium transients within SCEs over the total number of transients (left panel) and transient frequency (right panel), measured in P6-P8 mouse pups during the 0-10 epoch (N=12 animals), were compared with those described in a previous study (N=13 animals (Dard et al., 2022). Each dot represents one animal and the black line corresponds to the median. No difference was observed for the transient in SCEs ratio (Mann-Whitney test, U=55, p=0.2254) and for the transient frequency (Mann-Whitney test, U=57, p=0.2701). **(C)** FOV of the *stratum pyramidale* imaged during the 0-10 (left) and 40-50 (middle) periods from Emx1-Cre or VGluT2-Cre newborn mouse pups injected in the EC or VMT with mCherry or inhibitory DREADD virus. Scale bars = 100 µm. On the right, the contour map of imaged neurons from the FOV. Inferred *so*/*sp* GABAergic neurons in red (*so*/*sp*INs), inferred pyramidal cells in green (Pyr). **(D)** Raster plots showing neuronal activity as a function of time during the 0-10 (white) and 40-50 (light gray) periods of a newborn mouse pup injected with inhibitory mCherry virus. Graphs below represent the percentage of active cells as a function of time for both periods. **(E)** Same as **(D)** for animal stereotaxically injected at P0 with DREADD and subcutaneously with saline solution during calcium imaging experiment. **(F)** Cell transient frequency of mCherry/CNO (n=1615 cells, N=2 animals, left panel) and DREADD/Saline (n=2032 cells, N=3 animals, right panel) controls during 0-10 (light gray) and 40-50 (dark gray) epochs. Each boxplot shows the 25th, 50th and 75th percentiles and whiskers represent the 5th to the 95th percentile. The transient frequency was not different between the 0-10 and the 40-50 epochs for both mCherry/CNO (0-10: median=0.57 transient/min [iqr=8.90]; 40-50: median=0.57 transient/min [iqr=9.67]; Wilcoxon test, W=- 48137, p=0.1650) and DREADD/Saline (0-10: median=0.86 transient/min [iqr=15.22]; 40-50: median=0.96 transient/min [iqr=14.54]; Wilcoxon test, W=6717, p=0.8923). **(G)** Transients frequency measured during 40-50 period in *so*/*sp* GABAergic neurons (*so*/*sp*INs) and pyramidal cells (Pyr) normalized by 0-10 epoch transients frequency for mCherry/CNO (left) or DREADD/Saline (right) mice. Each plot shows the 50th and 95th percentile. For the mCherry/CNO, the median was not different from 100 for both *so*/*sp*INs (n=126 cells, median=100 % [iqr=1300], One sample Wilcoxon test, theoretical median 100, W=1233, p=0.1338) and Pyr (n=1299 cells, median=93.32 % [iqr=1500], One sample Wilcoxon test, theoretical median 100, W=-35702, p=0.1868). Similarly, for DREADD/Saline condition, there was no difference for for both *so*/*sp*INs (n=282 cells, median=94.85 % [iqr=5000], One sample Wilcoxon test, theoretical median 100, W=2601, p=0.3429) and Pyr (n=1498 cells, median=91.22 % [iqr=4000], One sample Wilcoxon test, theoretical median 100, W=30869, p=0.3567). **(H)** Animal behavior (number of movements, left panel; total movements duration, right panel; N= 9 animals) during the 0-10 and 40-50 epochs. Each black dot represents one animal. The number of movements was significantly different between 0-10 and 40-50 epochs (0-10: median= 55 movements [iqr=84]; 40-50: median= 71 movements [iqr=100]; Wilcoxon test, W=40, p=0.0156). The total movements duration was not different between 0-10 and 40-50 epochs (0-10: median=147.2 s [iqr=196.5]; 40-50: median=92.8 s [iqr=226.4]; Wilcoxon test, W=-17, p=0.3594).

**Figure S8.**
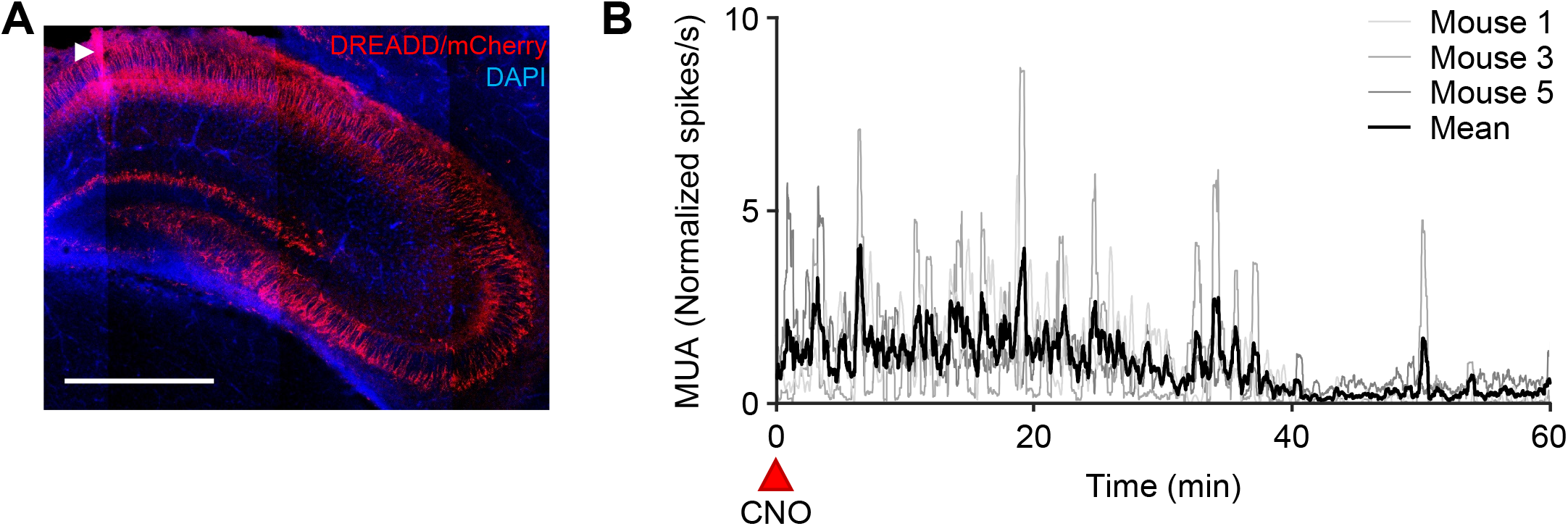
Control of CNO action time during *in vivo* extracellular electrophysiological recordings in mouse pups. **(A)** Example of an epifluorescence microscopy image of coronal slices of a P6-P8 brain of Emx1-Cre mice injected at P0 with DREADD/mCherry virus in the hippocampus, recorded with a silicon probe. The arrowhead shows the recording site of the probe stained with DiI. Scale bars: 500 µm. **(B)** Normalized CA1 multi-unit activity (MUA) of P6-P8 Emx1-Cre mouse pups injected in the hippocampus with the inhibitory DREADD virus, recorded for 1 hour following the subcutaneous injection of CNO. Each gray line represents an animal (n=3 animals), the dark line corresponds to the mean.

**Supplementary Movie 1. Light-sheet z-stack of a transparised brain from an Emx1-Cre mouse pup injected with Chronos/tdTomato virus in the EC.**

**Supplementary Movie 2. Light-sheet z-stack of a transparised brain from a VGluT2-Cre mouse pup injected with Chronos/tdTomato virus in the VMT.**

**Supplementary Movie 3. Example of an *ex vivo* calcium imaging movie from a P6 hippocampal slice from a wild-type mouse pup, injected with Chronos/tdTomato virus in the VMT at P0.** The white frames correspond to the light stimulation. The movie was sped up to ten times the acquisition rate.

**Supplementary Movie 4. Example of an *ex vivo* calcium imaging movie from a P6 hippocampal slice from a wild-type mouse pup, injected with Chronos/tdTomato virus in the EC at P0.** The white frames correspond to the light stimulation. The movie was sped up to ten times the acquisition rate.

**Supplementary Movie 5. Example of *in vivo* calcium imaging movies from P7 Emx1-Cre mouse pups injected with an inhibitory DREADD virus in the EC at P0.** The left and the right parts correspond, respectively, to the first ten minutes (0-10) and the fortieth to fiftieth minutes (40-50) after CNO injection. The movie was sped up to ten times the acquisition rate.

**Supplementary Movie 6. Example of *in vivo* calcium imaging movies from P8 VGluT2-Cre mouse pups injected with an inhibitory DREADD virus in the VMT at P0.** The left and the right parts correspond, respectively, to the first ten minutes (0-10) and the fortieth to fiftieth minutes (40-50) after CNO injection. The movie was sped up to ten times the acquisition rate.

**Supplementary Movie 7. Example of *in vivo* calcium imaging movies from a control P7 VGluT2-Cre mouse pups injected with an inhibitory DREADD virus in the VMT at P0.** The left and the right parts correspond, respectively, to the ten first minutes (0-10) and the fortieth to fiftieth minutes (40-50) after saline injection. The movie was sped up to ten times the acquisition rate.

**Supplementary Movie 8. Example of *in vivo* calcium imaging movies from a control P7 VGluT2-Cre mouse pups injected with a mCherry virus in the VMT at P0.** The left and the right parts correspond, respectively, to the ten first minutes (0-10) and the fortieth to fiftieth minutes (40-50) after CNO injection. The movie was sped up to ten times the acquisition rate.

## REFERENCES

Allene, C., Cattani, A., Ackman, J.B., Bonifazi, P., Aniksztejn, L., Ben-Ari, Y., and Cossart, R. (2008). Sequential Generation of Two Distinct Synapse-Driven Network Patterns in Developing Neocortex. Journal of Neuroscience 28, 12851–12863. https://doi.org/10.1523/jneurosci.3733-08.2008.

Andrianova, L., Brady, E.S., Margetts-Smith, G., Kohli, S., McBain, C.J., and Craig, M.T. (2021). Hippocampal CA1 pyramidal cells do not receive monosynaptic input from thalamic nucleus reuniens. Biorxiv 2021.09.30.462517. https://doi.org/10.1101/2021.09.30.462517.

Anstötz, M., Huang, H., Marchionni, I., Haumann, I., Maccaferri, G., and Lübke, J.H.R. (2015). Developmental Profile, Morphology, and Synaptic Connectivity of Cajal–Retzius Cells in the Postnatal Mouse Hippocampus. Cerebral Cortex 18, bhv271–18. https://doi.org/10.1093/cercor/bhv271.

Basu, J., Zaremba, J.D., Cheung, S.K., Hitti, F.L., Zemelman, B.V., Losonczy, A., and Siegelbaum, S.A. (2016). Gating of hippocampal activity, plasticity, and memory by entorhinal cortex long-range inhibition. Science 351, aaa5694–aaa5694. https://doi.org/10.1126/science.aaa5694.

Bayer, S.A. (1980). Development of the Hippocampal Region in the Rat I. Neurogenesis Examined With 3H-Thymidine Autoradiograph. The Journal of Comparative Neurology 190, 87–114.

Ben-Ari, Y., Cherubini, E., Corradetti, R., and Gaiarsa, J.L. (1989). Giant synaptic potentials in immature rat CA3 hippocampal neurones. The Journal of Physiology 416, 303–325.

Blankenship, A.G., and Feller, M.B. (2009). Mechanisms underlying spontaneous patterned activity in developing neural circuits. Nature Reviews Neuroscience 11, 18–29. https://doi.org/10.1038/nrn2759.

Bocchio, M., Gouny, C., Angulo-Garcia, D., Toulat, T., Tressard, T., Quiroli, E., Baude, A., and Cossart, R. (2020). Hippocampal hub neurons maintain distinct connectivity throughout their lifetime. Nature Communications 11, 1–19. https://doi.org/10.1038/s41467-020-18432-6.

Bokor, H., Csáki, Á., Kocsis, K., and Kiss, J. (2002). Cellular architecture of the nucleus reuniens thalami and its putative aspartatergic/glutamatergic projection to the hippocampus and medial septum in the rat. Eur J Neurosci 16, 1227–1239. https://doi.org/10.1046/j.1460-9568.2002.02189.x.

Bonifazi, P., Goldin, M., Picardo, M.A., Jorquera, I., Cattani, A., Bianconi, G., Represa, A., Ben-Ari, Y., and Cossart, R. (2009). GABAergic Hub Neurons Orchestrate Synchrony in Developing Hippocampal Networks. Science 326, 1419–1424. https://doi.org/10.1126/science.1175509.

Broussard, G.J., Liang, Y., Fridman, M., Unger, E.K., Meng, G., Xiao, X., Ji, N., Petreanu, L., and Tian, L. (2018). In vivo measurement of afferent activity with axon-specific calcium imaging. Nat Neurosci 21, 1272–1280. https://doi.org/10.1038/s41593-018-0211-4.

Cassel, J.-C., Ferraris, M., Quilichini, P., Cholvin, T., Boch, L., Stephan, A., and Vasconcelos, A.P. de (2021). The reuniens and rhomboid nuclei of the thalamus: a crossroads for cognition-relevant information processing? Neurosci Biobehav Rev 126, 338–360. https://doi.org/10.1016/j.neubiorev.2021.03.023.

Che, A., Babij, R., Iannone, A.F., Fetcho, R.N., Ferrer, M., Liston, C., Fishell, G., and García, N.V.D.M. (2018). Layer I Interneurons Sharpen Sensory Maps during Neonatal Development. Neuron 99, 98–116.e7. https://doi.org/10.1016/j.neuron.2018.06.002.

Chen, T.-W., Wardill, T.J., Sun, Y., Pulver, S.R., Renninger, S.L., Baohan, A., Schreiter, E.R., Kerr, R.A., Orger, M.B., Jayaraman, V., et al. (2013). Ultra-sensitive fluorescent proteins for imaging neuronal activity. Nature 499, 295–300. https://doi.org/10.1038/nature12354.

Chittajallu, R., Wester, J.C., Craig, M.T., Barksdale, E., Yuan, X.Q., Akgül, G., Fang, C., Collins, D., Hunt, S., Pelkey, K.A., et al. (2017). Afferent specific role of NMDA receptors for the circuit integration of hippocampal neurogliaform cells. Nature Communications 8, 152. https://doi.org/10.1038/s41467-017-00218-y.

Cossart, R., and Garel, S. (2022). Step by step: cells with multiple functions in cortical circuit assembly. Nat Rev Neurosci 1–16. https://doi.org/10.1038/s41583-022-00585-6.

Cossart, R., and Khazipov, R. (2021). How development sculpts hippocampal circuits and function. Physiol Rev https://doi.org/10.1152/physrev.00044.2020.

Crépel, V., Aronov, D., Jorquera, I., Represa, A., Ben-Ari, Y., and Cossart, R. (2007). A Parturition-Associated Nonsynaptic Coherent Activity Pattern in the Developing Hippocampus. Neuron 54, 105–120. https://doi.org/10.1016/j.neuron.2007.03.007.

Dard, R.F., Leprince, E., Denis, J., Rao-Balappa, S., Suchkov, D., Boyce, R., Lopez, C., Giorgi-Kurz, M., Szwagier, T., Dumont, T., et al. (2022). The rapid developmental rise of somatic inhibition disengages hippocampal dynamics from self-motion. Biorxiv 2021.06.08.447542. https://doi.org/10.1101/2021.06.08.447542.

Denis, J., Dard, R.F., Quiroli, E., Cossart, R., and Picardo, M.A. (2020). DeepCINAC: A Deep-Learning-Based Python Toolbox for Inferring Calcium Imaging Neuronal Activity Based on Movie Visualization. Eneuro 7, ENEURO.0038-20.2020. https://doi.org/10.1523/eneuro.0038-20.2020.

Donato, F., Jacobsen, R.I., Moser, M.-B., and Moser, E.I. (2017). Stellate cells drive maturation of the entorhinal-hippocampal circuit. Science eaai8178–17. https://doi.org/10.1126/science.aai8178.

Dzhala, V., Valeeva, G., Glykys, J., Khazipov, R., and Staley, K. (2012). Traumatic Alterations in GABA Signaling Disrupt Hippocampal Network Activity in the Developing Brain. Journal of Neuroscience 32, 4017–4031. https://doi.org/10.1523/jneurosci.5139-11.2012.

Ferraris, M., Cassel, J.-C., Vasconcelos, A.P. de, Stephan, A., and Quilichini, P.P. (2021). The Nucleus Reuniens, a thalamic relay for cortico-hippocampal interaction in recent and remote memory consolidation. Neurosci Biobehav Rev 125, 339–354. https://doi.org/10.1016/j.neubiorev.2021.02.025.

Franklin, K.B.J., and Paxinos, G. (2007). The Mouse Brain in Stereotaxic Coordinates, 3rd edition.

Gorski, J.A., Talley, T., Qiu, M., Puelles, L., Rubenstein, J.L.R., and Jones, K.R. (2002). Cortical Excitatory Neurons and Glia, But Not GABAergic Neurons, Are Produced in the Emx1-Expressing Lineage. J Neurosci 22, 6309–6314. https://doi.org/10.1523/jneurosci.22-15-06309.2002.

Hartung, H., Brockmann, M.D., Pöschel, B., Feo, V.D., and Hanganu-Opatz, I.L. (2016). Thalamic and Entorhinal Network Activity Differently Modulates the Functional Development of Prefrontal-Hippocampal Interactions. The Journal of Neuroscience : The Official Journal of the Society for Neuroscience 36, 3676–3690. https://doi.org/10.1523/jneurosci.3232-15.2016.

Hoover, W.B., and Vertes, R.P. (2012). Collateral projections from nucleus reuniens of thalamus to hippocampus and medial prefrontal cortex in the rat: a single and double retrograde fluorescent labeling study. Brain Struct Funct 217, 191–209. https://doi.org/10.1007/s00429-011-0345-6.

Ibrahim, L.A., Huang, S., Fernandez-Otero, M., Sherer, M., Qiu, Y., Vemuri, S., Xu, Q., Machold, R., Pouchelon, G., Rudy, B., et al. (2021). Bottom-up inputs are required for establishment of top-down connectivity onto cortical layer 1 neurogliaform cells. Neuron https://doi.org/10.1016/j.neuron.2021.08.004.

Karlsson, K.A.E., Mohns, E.J., Prisco, G.V. di, and Blumberg, M.S. (2006). On the co-occurrence of startles and hippocampal sharp waves in newborn rats. Hippocampus 16, 959–965. https://doi.org/10.1002/hipo.20224.

Kerr, K.M., Agster, K.L., Furtak, S.C., and Burwell, R.D. (2007). Functional neuroanatomy of the parahippocampal region: The lateral and medial entorhinal areas. Hippocampus 17, 697–708. https://doi.org/10.1002/hipo.20315.

Kim, J.-Y., Grunke, S.D., Levites, Y., Golde, T.E., and Jankowsky, J.L. (2014). Intracerebroventricular Viral Injection of the Neonatal Mouse Brain for Persistent and Widespread Neuronal Transduction. J Vis Exp 51863. https://doi.org/10.3791/51863.

Klapoetke, N.C., Murata, Y., Kim, S.S., Pulver, S.R., Birdsey-Benson, A., Cho, Y.K., Morimoto, T.K., Chuong, A.S., Carpenter, E.J., Tian, Z., et al. (2014). Independent Optical Excitation of Distinct Neural Populations. Nat Methods 11, 338–346. https://doi.org/10.1038/nmeth.2836.

Krashes, M.J., Koda, S., Ye, C., Rogan, S.C., Adams, A.C., Cusher, D.S., Maratos-Flier, E., Roth, B.L., and Lowell, B.B. (2011). Rapid, reversible activation of AgRP neurons drives feeding behavior in mice. J Clin Invest 121, 1424–1428. https://doi.org/10.1172/jci46229.

Luhmann, H.J., and Khazipov, R. (2017). Neuronal activity patterns in the developing barrel cortex. Neuroscience 368, 256–267. https://doi.org/10.1016/j.neuroscience.2017.05.025.

Marques-Smith, A., Lyngholm, D., Kaufmann, A.-K., Stacey, J.A., Hoerder-Suabedissen, A., Becker, E.B.E., Wilson, M.C., Molnár, Z., and Butt, S.J.B. (2016). A Transient Translaminar GABAergic Interneuron Circuit Connects Thalamocortical Recipient Layers in Neonatal Somatosensory Cortex. Neuron 89, 536–549. https://doi.org/10.1016/j.neuron.2016.01.015.

Martini, F.J., Guillamón-Vivancos, T., Moreno-Juan, V., Valdeolmillos, M., and López-Bendito, G. (2021). Spontaneous activity in developing thalamic and cortical sensory networks. Neuron 109, 2519–2534. https://doi.org/10.1016/j.neuron.2021.06.026.

Masurkar, A.V., Srinivas, K.V., Brann, D.H., Warren, R., Lowes, D.C., and Siegelbaum, S.A. (2017). Medial and Lateral Entorhinal Cortex Differentially Excite Deep versus Superficial CA1 Pyramidal Neurons. CellReports 18, 148–160. https://doi.org/10.1016/j.celrep.2016.12.012.

Masurkar, A.V., Tian, C., Warren, R., Reyes, I., Lowes, D.C., Brann, D.H., and Siegelbaum, S.A. (2020). Postsynaptic integrative properties of dorsal CA1 pyramidal neuron subpopulations. Journal of Neurophysiology 123, 980–992. https://doi.org/10.1152/jn.00397.2019.

Melzer, S., Michael, M., Caputi, A., Eliava, M., Fuchs, E.C., Whittington, M.A., and Monyer, H. (2012). Long-Range-Projecting GABAergic Neurons Modulate Inhibition in Hippocampus and Entorhinal Cortex. Science 335, 1506–1510. https://doi.org/10.1126/science.1217139.

Minlebaev, M., Colonnese, M., Tsintsadze, T., Sirota, A., and Khazipov, R. (2011). Early γ oscillations synchronize developing thalamus and cortex. Science 334, 226–229. https://doi.org/10.1126/science.1210574.

Mizuno, H., Ikezoe, K., Nakazawa, S., Sato, T., Kitamura, K., and Iwasato, T. (2018). Patchwork-Type Spontaneous Activity in Neonatal Barrel Cortex Layer 4 Transmitted via Thalamocortical Projections. CellReports 22, 123–135. https://doi.org/10.1016/j.celrep.2017.12.012.

Mohns, E.J., and Blumberg, M.S. (2008). Synchronous Bursts of Neuronal Activity in the Developing Hippocampus: Modulation by Active Sleep and Association with Emerging Gamma and Theta Rhythms. Journal of Neuroscience 28, 10134–10144. https://doi.org/10.1523/jneurosci.1967-08.2008.

Molnár, Z., Luhmann, H.J., and Kanold, P.O. (2020). Transient cortical circuits match spontaneous and sensory-driven activity during development. Science 370, eabb2153. https://doi.org/10.1126/science.abb2153.

Murata, Y., and Colonnese, M.T. (2020). GABAergic interneurons excite neonatal hippocampus in vivo. Science Advances 6, eaba1430. https://doi.org/10.1126/sciadv.aba1430.

Nakagawa, Y. (2019). Development of the thalamus: From early patterning to regulation of cortical functions. Wiley Interdiscip Rev Dev Biology 8, e345. https://doi.org/10.1002/wdev.345.

Overstreet-Wadiche, L., and McBain, C.J. (2015). Neurogliaform cells in cortical circuits. Nat Rev Neurosci 16, 458–468. https://doi.org/10.1038/nrn3969.

Pachitariu, M., Stringer, C., Dipoppa, M., Schröder, S., Rossi, L.F., Dalgleish, H., Carandini, M., and Harris, K.D. (2017). Suite2p: beyond 10,000 neurons with standard two-photon microscopy. Biorxiv 061507. https://doi.org/10.1101/061507.

Paxinos, Halliday, Watson, and Kassem (2020). Atlas of the Developing Mouse Brain.

Pnevmatikakis, E.A., and Giovannucci, A. (2017). NoRMCorre: An online algorithm for piecewise rigid motion correction of calcium imaging data. Journal of Neuroscience Methods 291, 83–94. https://doi.org/10.1016/j.jneumeth.2017.07.031.

Quattrocolo, G., and Maccaferri, G. (2014). Optogenetic Activation of Cajal-Retzius Cells Reveals Their Glutamatergic Output and a Novel Feedforward Circuit in the Developing Mouse Hippocampus. J Neurosci 34, 13018–13032. https://doi.org/10.1523/jneurosci.1407-14.2014.

Rio-Bermudez, C.D., and Blumberg, M.S. (2021). Sleep as a window on the sensorimotor foundations of the developing hippocampus. Hippocampus https://doi.org/10.1002/hipo.23334.

Ronzitti, E., Conti, R., Zampini, V., Tanese, D., Foust, A.J., Klapoetke, N., Boyden, E.S., Papagiakoumou, E., and Emiliani, V. (2017). Submillisecond Optogenetic Control of Neuronal Firing with Two-Photon Holographic Photoactivation of Chronos. The Journal of Neuroscience : The Official Journal of the Society for Neuroscience 37, 10679–10689. https://doi.org/10.1523/jneurosci.1246-17.2017.

Roth, B.L. (2016). DREADDs for Neuroscientists. Neuron 89, 683–694. https://doi.org/10.1016/j.neuron.2016.01.040.

Rübel, O., Tritt, A., Ly, R., Dichter, B.K., Ghosh, S., Niu, L., Soltesz, I., Svoboda, K., Frank, L., and Bouchard, K.E. (2022). The Neurodata Without Borders ecosystem for neurophysiological data science. Biorxiv 2021.03.13.435173. https://doi.org/10.1101/2021.03.13.435173.

Siegel, F., Heimel, J.A., Peters, J., and Lohmann, C. (2012). Peripheral and Central Inputs Shape Network Dynamics in the Developing Visual Cortex In Vivo. Current Biology 22, 253–258. https://doi.org/10.1016/j.cub.2011.12.026.

Siegle, J.H., López, A.C., Patel, Y.A., Abramov, K., Ohayon, S., and Voigts, J. (2017). Open Ephys: an open-source, plugin-based platform for multichannel electrophysiology. J Neural Eng 14, 045003. https://doi.org/10.1088/1741-2552/aa5eea.

Supèr, H., and Soriano, E. (1994). The organization of the embryonic and early postnatal murine hippocampus. II. Development of entorhinal, commissural, and septal connections studied with the lipophilic tracer DiI. The Journal of Comparative Neurology 344, 101–120. https://doi.org/10.1002/cne.903440108.

Tricoire, L., Pelkey, K.A., Daw, M.I., Sousa, V.H., Miyoshi, G., Jeffries, B., Cauli, B., Fishell, G., and McBain, C.J. (2010). Common Origins of Hippocampal Ivy and Nitric Oxide Synthase Expressing Neurogliaform Cells. Journal of Neuroscience 30, 2165–2176. https://doi.org/10.1523/jneurosci.5123-09.2010.

Tuncdemir, S.N., Wamsley, B., Stam, F.J., Osakada, F., Goulding, M., Callaway, E.M., Rudy, B., and Fishell, G. (2016). Early Somatostatin Interneuron Connectivity Mediates the Maturation of Deep Layer Cortical Circuits. Neuron 89, 521–535. https://doi.org/10.1016/j.neuron.2015.11.020.

Tyzio, R., Represa, A., Jorquera, I., Ben-Ari, Y., Gozlan, H., and Aniksztejn, L. (1999). The Establishment of GABAergic and Glutamatergic Synapses on CA1 Pyramidal Neurons is Sequential and Correlates with the Development of the Apical Dendrite. 1–11.

Valeeva, G., Tressard, T., Mukhtarov, M., Baude, A., and Khazipov, R. (2016). An Optogenetic Approach for Investigation of Excitatory and Inhibitory Network GABA Actions in Mice Expressing Channelrhodopsin-2 in GABAergic Neurons. J Neurosci Official J Soc Neurosci 36, 5961–5973. https://doi.org/10.1523/jneurosci.3482-15.2016.

Valeeva, G., Janackova, S., Nasretdinov, A., Rychkova, V., Makarov, R., Holmes, G.L., Khazipov, R., and Lenck-Santini, P.-P. (2019). Emergence of Coordinated Activity in the Developing Entorhinal–Hippocampal Network. Cereb Cortex 29, 906–920. https://doi.org/10.1093/cercor/bhy309.

Vertes, R.P. (2015). Chapter 7 Major diencephalic inputs to the hippocampus supramammillary nucleus and nucleus reuniens. Circuitry and function. Prog Brain Res 219, 121–144. https://doi.org/10.1016/bs.pbr.2015.03.008.

Villette, V., Malvache, A., Tressard, T., Dupuy, N., and Cossart, R. (2015). Internally Recurring Hippocampal Sequences as a Population Template of Spatiotemporal Information. Neuron 88, 357–366. https://doi.org/10.1016/j.neuron.2015.09.052.

Vong, L., Ye, C., Yang, Z., Choi, B., Chua, S., and Lowell, B.B. (2011). Leptin Action on GABAergic Neurons Prevents Obesity and Reduces Inhibitory Tone to POMC Neurons. Neuron 71, 142–154. https://doi.org/10.1016/j.neuron.2011.05.028.

Wang, Y., Romani, S., Lustig, B., Leonardo, A., and Pastalkova, E. (2015). Theta sequences are essential for internally generated hippocampal firing fields. Nature Neuroscience 18, 282–288. https://doi.org/10.1038/nn.3904.

Weel, M.J.D. der, and Witter, M.P. (2020). THE THALAMIC MIDLINE NUCLEUS REUNIENS: POTENTIAL RELEVANCE FOR SCHIZOPHRENIA AND EPILEPSY. Neurosci Biobehav Rev 119, 422–439. https://doi.org/10.1016/j.neubiorev.2020.09.033.

Weel, M.J.D.D., and Witter, M.P. (1996). Projections from the nucleus reuniens thalami to the entorhinal cortex, hippocampal field CA1, and the subiculum in the rat arise from different populations of neurons. J Comp Neurol 364, 637–650. https://doi.org/10.1002/(sici)1096-9861(19960122)364:4<637::aid-cne3>3.0.co;2-4.

Wester, J.C., and McBain, C.J. (2016). Interneurons Differentially Contribute to Spontaneous Network Activity in the Developing Hippocampus Dependent on Their Embryonic Lineage. The Journal of Neuroscience : The Official Journal of the Society for Neuroscience 36, 2646–2662. https://doi.org/10.1523/jneurosci.4000-15.2016.

Zhang, G.-W., Sun, W.-J., Zingg, B., Shen, L., He, J., Xiong, Y., Tao, H.W., and Zhang, L.I. (2018). A Non-canonical Reticular-Limbic Central Auditory Pathway via Medial Septum Contributes to Fear Conditioning. Neuron 97, 406–417.e4. https://doi.org/10.1016/j.neuron.2017.12.010.

Zsiros, V., Aradi, I., and Maccaferri, G. (2007). Propagation of postsynaptic currents and potentials via gap junctions in GABAergic networks of the rat hippocampus. J Physiology 578, 527–544. https://doi.org/10.1113/jphysiol.2006.123463.

Zutshi, I., Valero, M., Fernández-Ruiz, A., and Buzsáki, G. (2021). Extrinsic control and intrinsic computation in the hippocampal CA1 circuit. Neuron https://doi.org/10.1016/j.neuron.2021.11.015.

